# TNF Receptor 1 regulates colonic mesenchymal cell diversity and the epithelial stem cell niche

**DOI:** 10.1101/2025.09.08.674957

**Authors:** Safina Gadeock, Nandini Girish, Cambrian Y. Liu, Ying Huang, Tracy C. Grikscheit, D. Brent Polk

## Abstract

**BACKGROUND & AIMS:** Anti-tumor necrosis factor (anti-TNF) is a mainstay of inflammatory bowel disease (IBD) therapy but fails in many patients. Although TNF has pro-inflammatory effects, depletion of TNF receptor 1 (TNFR1) paradoxically exacerbates chronic colitis. Because colitis induces remodeling of mesenchymal cell populations, which provide a niche for epithelial stem cells involved in mucosal healing, we hypothesized that TNFR1 promotes colonic mesenchymal cell diversity and stem cell niche function.

**METHODS:** Mesenchymal TNFR1 function was studied using TNFR1^-/-^, platelet derived growth factor receptor alpha (PDGFRα)-Cre;TNFR1^fl/fl^ mice, and mixed-genotype mesenchymal-epithelial co-cultures. Mesenchymal cell diversity and gene function were assessed using single-cell RNA-Seq of primary colonic myofibroblasts (CMFs) and via anti-integrin A6 (ITGA6) antibody treatment and exogenous R-spondin 3 (RSPO3) supplementation.

**RESULTS:** TNFR1^-/-^ mesenchyme exhibits reduced cell diversity, with specific depletion of specialized TNF- and interferon-signaling pericryptal cell-type. Deletion of TNFR1 in the pericryptal mesenchyme diminished the (PDGFRα)+ CMF population and reduced RSPO3 expression, but increased ITGA6 expression relative to controls (TNFR1^+/-^). Moreover, inhibition of ITGA6 reversed the proliferative and migratory phenotype of TNFR1^-/-^ CMFs and restored expression of PDGFRα and RSPO3. Co-cultures of colonoids with TNFR1^-/-^ CMFs resulted in downregulation of stem cell marker expression; this was rescued by supplementation with RSPO3. Supporting the role for mesenchymal TNFR1 in regulating colonic epithelial stem cells, mice deficient for TNFR1 in PDGFRα+ cells showed a 40% loss of Lgr5+ stem cells, consistent with the global TNFR1-deficient mouse.

**CONCLUSION:** TNFR1-mediated signaling regulates specification and function of colonic mesenchyme, performing an integral role in the maintenance of the crypt stem cell population.

## Introduction

Inflammatory bowel disease (IBD) is associated with prominent increases in circulating and mucosal levels of tumor necrosis factor (TNF). ^1^ However, TNF’s selective effects on specific tissues in the intestinal mucosa are not fully understood. TNF binds two receptors, tumor necrosis factor receptor (TNFR) 1 (p55, *Tnfrsf1a*) and 2 (p75, *Tnfrsf1b*). TNFR1 exhibits widespread intestinal expression, including in epithelium and stroma, whereas TNFR2 is expressed specifically on immune cell subsets and is upregulated in epithelial cells in IBD. ^2, 3^ TNF is pathogenic in IBD as shown by initial successful treatment of the majority of patients with ulcerative colitis or Crohn’s disease. ^4^ TNF initiates pro-inflammatory signaling that leads to autoimmune tissue destruction and cellular apoptosis. ^5–7^ Paradoxically, TNF can also activate pathways to mediate cell survival, ^8^ epithelial regeneration, ^9, 10^ and immunological tolerance limiting colonic inflammation. ^11^ Thus, the pleiotropic biological effects of TNF, ^2, 12^ the exacerbation of colitis in multiple models of TNF pathway ablation, ^12, 13^ and the clinical data suggesting that most patients eventually fail anti-TNF therapy^4^ do not lend to a simple paradigm for TNF’s role in IBD. Deciphering the functions of TNF in the body thus remains an urgent clinical priority.

Mesenchymal stromal cells have been implicated in several processes central to mucosal healing^14^ and IBD-associated inflammation. ^15^ The colonic myofibroblast (CMF) population, which harbors a specialized pericryptal mesenchymal subpopulation, provides critical molecular cues (e.g., Wnt ligands, transforming growth factors, insulin growth factors, Notch ligands, epidermal growth factor, and Bmp-type molecules) that maintain the self-renewal of crypt stem cells and guide the proper differentiation of epithelial cells along the crypt vertical axis. ^14, 16^ During epithelial wound healing, mesenchymal cells secrete noncanonical Wnt ligands that drive crypt morphogenesis and cell proliferation. ^17, 18^ Thus, the function of mesenchymal cells is intimately associated with epithelial regenerative capacity in homeostasis and during states of challenge. Whether cytokines associated with IBD might regulate mesenchymal behaviors themselves, and consequently the epithelial stem cell pool, remains to be determined, as it is primarily considered that cytokines act cell-autonomously on epithelium. In fact, recent single-cell profiling has demonstrated that the colonic mesenchyme undergoes significant remodeling in IBD, ^15, 19–21^ which would suggest that the inflammatory milieu has a profound ability to regulate specification and function of CMFs. Here, we tested the hypothesis that TNF signaling through TNFR1 regulates the diversity, function, and niche-forming abilities of colonic CMFs. By studying the receptor rather than the ligand, we could target the analysis specifically to CMFs. The reported results link TNF/TNFR1 signaling to maintenance of the epithelial stem cell pool through the profound effects of this cytokine on the mesenchymal “support” cells of the colonic crypt.

## Results

### Mesenchymal TNFR1 deletion alters colonic mesenchymal marker expression

TNFR1 promotes intestinal epithelial cell survival, ^8^ stem cell proliferation, ^10^ colonic barrier integrity, mucosal immune cell function, and repair pathways to restrict IBD-associated inflammation. ^11^ Its role in the intestinal mesenchyme, however, remains unknown. To characterize the role of TNFR1 in mesenchymal cell specification and function, we first compared mesenchymal cell populations in TNFR1^-/-^ and control (TNFR1^+/-^) mice. There are several known and potentially overlapping colonic mesenchymal cell populations, which are defined by expression of alpha-smooth muscle actin (αSMA), vimentin (VIM), platelet derived growth factor receptor alpha (PDGFRα), GLI family zinc finger 1 (GLI1), and/or forkhead box L1 (FOXL1). We found a 40% reduction in VIM but a 58% increase in αSMA expression in TNFR1^-/-^ mice (Figures 1*AI*, 1*AII*, 1*BI*, and 1*BII*). Focusing on the pericryptal cells previously shown to serve as a stem cell niche, ^22^ we quantified the total levels of FOXL1, GLI1 and PDGFRα+ mesenchymal populations. The expression of PDGFRα was decreased by 30% in colons obtained from TNFR1^-/-^ mice (Figures 1*AIII* and 1*BIII*). However, the expressions of GLI1 or FOXL1 were unchanged (Figures 1*AIV*, 1*AV*, 1*BIV*, and 1*BV*).

**Figure 1.**
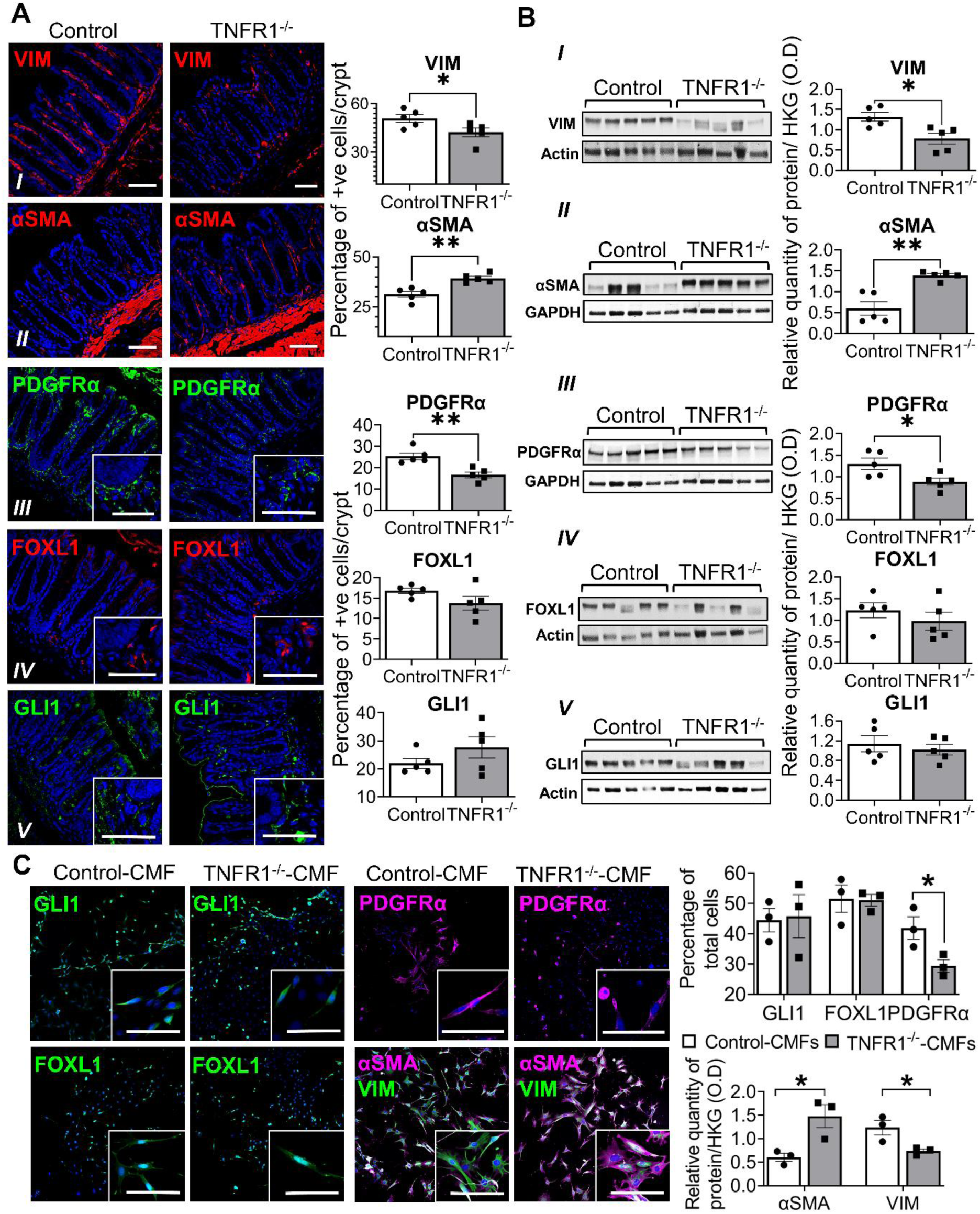
Mesenchymal TNFR1 deletion alters colonic mesenchymal marker expression. (A) Immunolocalization and percentage of mesenchymal cell markers, and (B) total proteins levels of (*I*) vimentin, (*II*) αSMA, (*III*) PDGFRα, (*IV*) FOXL1, and (*V*) GLI1 in sub-epithelial MSCs of the distal colon of 16wk-old control (TNFR1^+/-^) and TNFR1^-/-^ mice. Subsets represent magnified localization of the markers in the pericryptal niche. (C) Immunolocalization and quantification of GLI1, FOXL1, PDGFRα+ cells, and αSMA and vimentin expression in control or TNFR1^-/-^ CMFs. Scale bars = 50µm. *N* = 3-5 animals. Error bars are presented as mean ± SEM. Data are analyzed by an unpaired Student *t* test, where * P < 0.05; ** P < 0.01, and ^#^ P = 0.05-0.09.

To determine if the specific expression of TNFR1 on the sub-epithelial mesenchyme was associated with altered specification of mesenchymal populations, we generated Pdgfra-Cre;TNFR1^fl/fl^ animals (Supplementary Figures 1AI and 1*AII*) to ablate TNFR1 in sub-epithelial PDGFRα+ cells. We show that while αSMA+ cells increased by 40%; pericryptal PDGFRα+ cells were downregulated by 25% in Pdgfra-Cre;TNFR1^fl/fl^ animals relative to TNFR1^fl/fl^ and Pdgfra-Cre controls (Supplementary Figures 1B and 1C). Epithelial-(Vil1-Cre;TNFR1^fl/fl^) or myeloid cell-specific (LysM-Cre;TNFR1^fl/fl^) deletion of TNFR1 did not affect the expression of mesenchymal cell markers (Supplementary Figures 2A and 2B). We have shown previously that intestinal mucosal ablation of TNFR1 results in an altered phenotype resulting in increased neutrophils accumulation, loss of epithelial proliferation, increase in DNA-damage markers and crypt abscess.^11^ Here, we show that this phenotype is associated with an altered mesenchymal lineage in TNFR1^-/-^ animals relative to TNFR1^+/-^ controls. We also demonstrate that a similar phenotype of increased neutrophils accumulation, loss of epithelial proliferation, increase in DNA-damage markers is preserved in Pdgfra-Cre;TNFR1^fl/fl^ animals relative to TNFR1^fl/fl^ and Pdgfra-Cre controls (Supplementary Figures 3A-3C), Thus, cell-autonomous TNFR1 deletion in PDGFRα+ cells *in vivo* selectively alters the colonic mesenchymal cell census.

We next asked if we could recover key mesenchymal subpopulations *in vitro*. After 5 days of three-dimensional culture, we recovered partially overlapping CMF populations that were VIM+, αSMA+, PDGFRα+, GLI1+, or FOXL1+ (Figure 1*C* and Supplementary Figures 4A, and 4*B*). These populations were negative for desmin, a marker of muscle cells. Crucially, and consistent with our observations in TNFR1^-/-^ mice *in vivo*, the proportion of the cultured PDGFRα+ mesenchymal subpopulation within CMFs was reduced by 21% in TNFR1^-/-^ conditions (Figures 1*C* and Supplementary Figure 4B). Western blot analysis of mesenchymal cell homogenates showed increased αSMA expression and decreased vimentin expression in TNFR1^-/-^ CMFs (Figure 1*C*). Therefore, culture conditions recapitulate key features of the role of TNFR1 in maintaining homeostatic representation of CMFs.

### TNFR1 is required to preserve colonic mesenchymal cell diversity including a novel subpopulation defined by TNF and IFN signaling genes

Significant alterations in markers of mesenchymal cells in TNFR1^-/-^ mice and *in vitro* suggest changes in overall CMF diversity. To test this prediction, we performed single-cell RNA sequencing of murine mesenchymal cells, isolated and amplified from 3 independent TNFR1^+/-^ (littermate controls) and 3 TNFR1^-/-^ mice *in vitro*. The cultures were prepared for sequencing concurrently and processed as a single batch. In total, we recovered >34,000 cells (Figure 2*A*) from the 6 samples. Clustering of cells comprising the merged dataset revealed 6 main clusters (Figure 2*B*). Notably, clusters 1 and 4 were largely linked to cultures derived from TNFR1^-/-^ mice, with limited overlap in markers seen in the TNFR1^+/-^ controls (Figures 2*B* and 2*C*). In contrast, clusters 2, 3, 5, and 6 were predominantly associated with cultures from control mice (Figure 2*B* and 2*C*). This distribution was consistent across all cultures analyzed. Together, these findings indicate that loss of TNFR1 leads to an almost complete inability of CMFs to adopt homeostatic cellular phenotypes, reflecting a defect in CMF cell-type specification.

**Figure 2.**
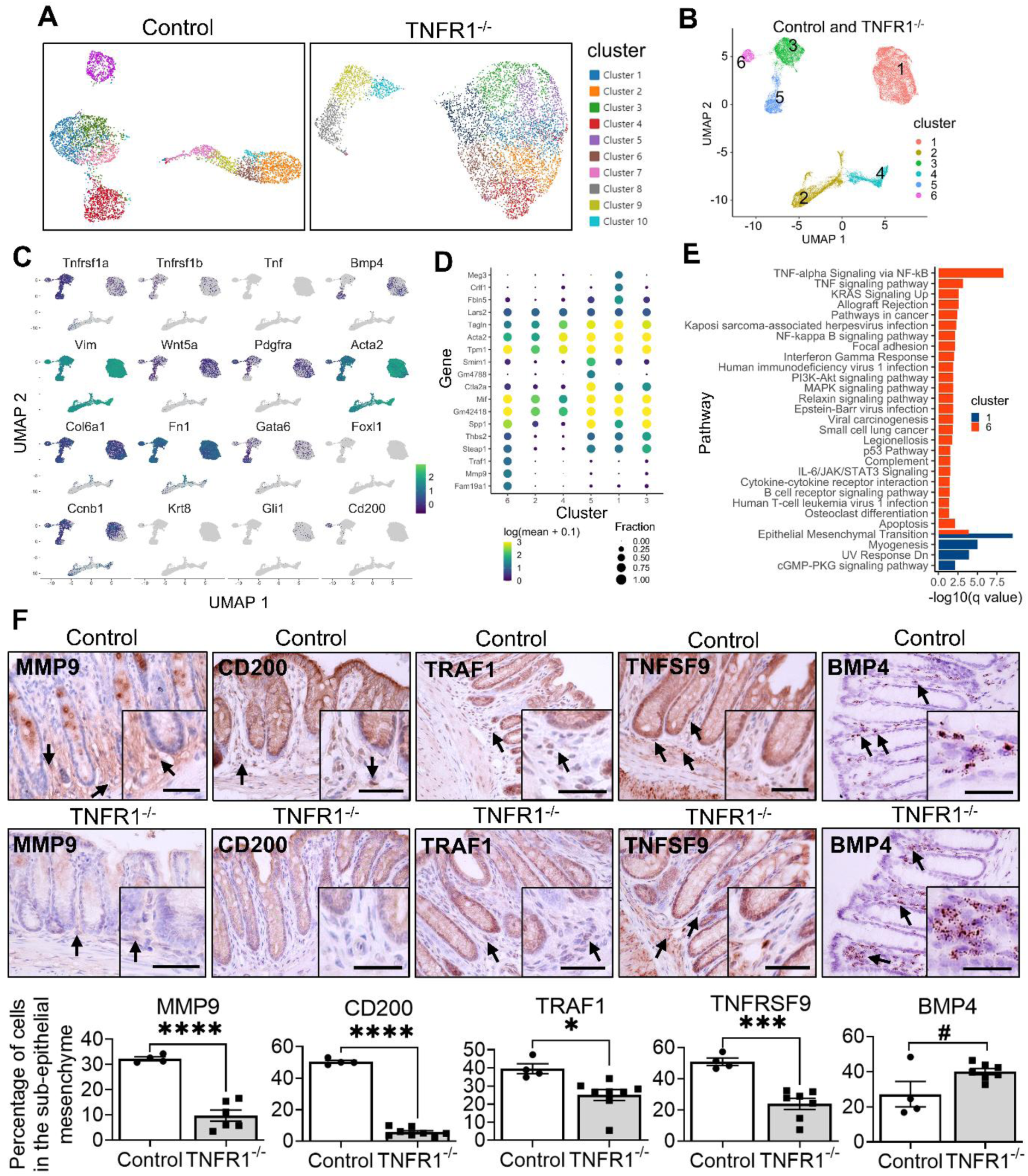
TNFR1 is required to preserve colonic mesenchymal cell diversity including a novel subpopulation defined by TNF and IFN signaling genes. (A) UMAP identifies 10 clusters in unsupervised raw counts from primary CMFs derived from control (TNFR1^+/-^) and TNFR1^-/-^ mice showing the differential cell populations in each genotype. (*N* = 3) (B) UMAP identifies 6 clusters in 34,415 primary CMFs derived from control (TNFR1^+/-^) and TNFR1^-/-^ mice, merged and clustered together after sequencing, reflecting completely individualistic clusters. (C) Expression of classical mesenchymal identity markers in control and TNFR1^-/-^ CMFs, segregated across the clusters identified by scRNA-Seq. (D) Dot plot showing expression of significantly modified genes against detected clusters. (E) Gene ontology pathways showing significant enrichment among top marker genes for cluster 1 and 6 derived from control and TNFR1^-/-^ CMFs. (F) Immunostaining of MMP9, CD200, TNFRSF9, TRAF1, and BMP4 highly enriched in cluster 6 from control CMFs and significantly diminished in cluster 1 of TNFR1^-/-^ CMFs, in native colon of control and TNFR1^-/-^ mice. Black arrows indicate positively stained cells. Scale bars = 50µm. All images are representative of *N* = 4 – 8 animals. Error bars are presented as mean ± SEM. Data are analyzed by an unpaired Student *t* test, where * P < 0.05; *** P < 0.001 and **** P < 0.0001, and ^#^ P = 0.05-0.09.

We next mapped out the expression of known mesenchymal cell markers across the identified cell clusters. No *Krt8*+ epithelial cells were found in the data (Figure 2*C*), consistent with the purity of cultures demonstrated in Supplementary Figure 4A. Clusters 1, 3, 5, and 6 expressed the main TNF receptors, *Tnfrsf1a* and *Tnfrsf1b*, with cluster 1 (obtained from TNFR1^-/-^ mice) showing reduced levels of *Tnfrsf1a* expression (Figure 2*C*). All clusters expressed high levels of *Vim*, consistent with their mesenchymal identity (Figure 2*C*). Each cluster included a subpopulation of *Ccnb1*+ cells, indicating proliferative potential. Clusters 2 (control) and 4 (knockout) did not express any characteristic markers, but expressed general mesenchymal transcripts (e.g., *Acta2*, *Vim*) at high levels. Although the cells in these clusters passed quality inspection, we excluded them from further analysis. Clusters 1 (knockout), 3 (control), 5 (control), and 6 (control) expressed markers of CMFs, including *Pdgfra*, *Gata6*, *Col6a1*, *Fn1*, *Bmp4*, *Wnt5a*, *Gli1*, and *Foxl1*, suggesting that each cluster represents distinct subpopulations of CMFs. The most salient of these clusters was cluster 6, which expressed many unique markers including *Mmp9*, *Cd200*, *Tnfrsf9*, and *Traf1* that were depleted in TNFR1^-/-^ cultures (Figures 2*C* and 2*D*). Particularly, MMP9+ subepithelial mesenchymal cells were reduced by 25% in Pdgfra-Cre;TNFR1^fl/fl^ mice relative to TNFR1^fl/fl^ and Pdgfra-Cre controls (Supplementary Figure 1C). Moreover, their levels remained unchanged in the colonic mesenchyme of Vil1-Cre;TNFR1^fl/fl^ and LysM-Cre;TNFR1^fl/fl^ animals relative to TNFR1^fl/fl^ controls (Supplementary Figure 2C), demonstrating the specific role of pericryptal mesenchymal TNFR1 in the regulation of the cluster 6 cells.

TNFR1^-/-^ CMFs do not stratify into distinct cell-types. Thus, even though the recovered cell number in cluster 1 (knockout) was roughly equivalent to the sum of the cells in clusters 3, 5, and 6 (controls), the transcriptional diversity within the TNFR1^-/-^ CMF population was sufficiently reduced to preclude assignment of these knockout cells to more than one cluster. In contrast, there were 3 clusters of control CMFs.

To determine how these clusters relate to one another, we identified markers for each and categorized these markers according to representative signaling pathways. Gene markers of cluster 1 (knockout) included *Meg3*, *Crlf1*, and *Fbln5*; these 190 markers were related to epithelial-mesenchymal transition, myogenesis, downregulation of the UV response, and cGMP signaling (Figures 2*C* and 2*D*). Cluster 3 (control) was defined by 18 marker genes and was notable for its loss in Bmp4 expression and shared expression of *Col6a1* and *Fn1* with clusters 1 and 6 (Figures 2*C* and 2*D*); there were too few markers to identify enriched pathways. Cluster 5 (control) was specified by 293 markers and exhibited high expression of *Smim1* and *Ctla2a* (Figure 2*D*), but unexpectedly no significant pathways (q < 0.05) were identified. Cluster 6 also expressed many unique genes (190), including *Mmp9*, *Cd200* and *Traf1* (Figure 2*C* and 2*D*), and these were related to TNF signaling and interferon and other cytokine signaling (Figures 2*E* and Supplementary Figure 5). The loss of distinct cluster 6 markers in TNFR1^-/-^ CMFs, identified by UMAP and gene clustering in Figures 2*C* and 2*D*, was confirmed *in vivo* in TNFR1^-/-^ animals. These markers were selected based on their high magnitude of upregulation in cluster 6 of control CMFs, and their near-complete depletion in TNFR1^-/-^ CMFs. By immunostaining, there was a 70% reduction in MMP9+; 89% in CD200+; 37% in TRAF1+; and 53% in TNFRSF9+ cells; but differences in the numbers of BMP4+ cells in TNFR1^-/-^ animals relative to controls did not reach statistical significance (Figure 2*F*). Thus, cluster 6 represents a specialized population of mesenchymal cells distinguished by enrichment of TNF signaling pathways. In the absence of TNFR1, this cluster fails to emerge, and the CMF population is correspondingly less diverse. Thus, TNFR1 regulates the colonic homeostatic mesenchymal cell census and its diversification into specialized subpopulations. Crucially, it determines the presence of sub-epithelial mesenchymal cells that are a hub for TNF and cytokine signaling.

### Transcriptome profiling of TNFR1-deficient CMFs identifies upregulation of integrin A6 but loss of the epithelial stem cell niche factor R-spondin 3

Although single-cell experiments enable profiling of mesenchymal cell diversity, they can miss crucial gene products expressed at low levels that distinguish TNFR1^-/-^ from control CMFs. We therefore investigated expression of TNFR1^-/-^ compared to control CMFs via bulk RNA-Seq analysis of cultured primary cells. We found concordance between the bulk and single-cell transcriptomic approaches (Supplementary Figures 6A-6E). KEGG pathway analysis of differentially expressed genes showed that the loss of TNFR1 alters anchoring and cell substrate junctions, focal adhesions, extracellular matrix, and actin binding processes (Figure 3*A*). Related to enhanced matrix signaling, *Itga6*, representing the integrin alpha 6 subunit, was significantly upregulated in TNFR1^-/-^ compared to control CMFs (Figure 3*B*). We found a specific reduction in key stem cell niche factors such as *Rspo3*, *Wnt2b*, and *Fgf2* in knockout CMFs (Figure 3*C*). In particular, *Rspo3*, crucial for Lgr5+ stem cell maintenance, ^22, 23^ was downregulated 5.1-fold in TNFR1^-/-^ CMFs (Figure 3*C* and 3*D*). *In vivo*, we found that in Pdgfra-Cre;TNFR1^fl/fl^ animals, Rspo3 levels were reduced in the pericryptal mesenchyme (Supplementary Figure 1C). This effect was not observed with epithelial- or myeloid-specific deletion of TNFR1 (Supplementary Figure 2C). Thus, pericryptal mesenchymal TNFR1 regulates expression of the Rspo3 niche factor in cell-autonomous manner.

**Figure 3.**
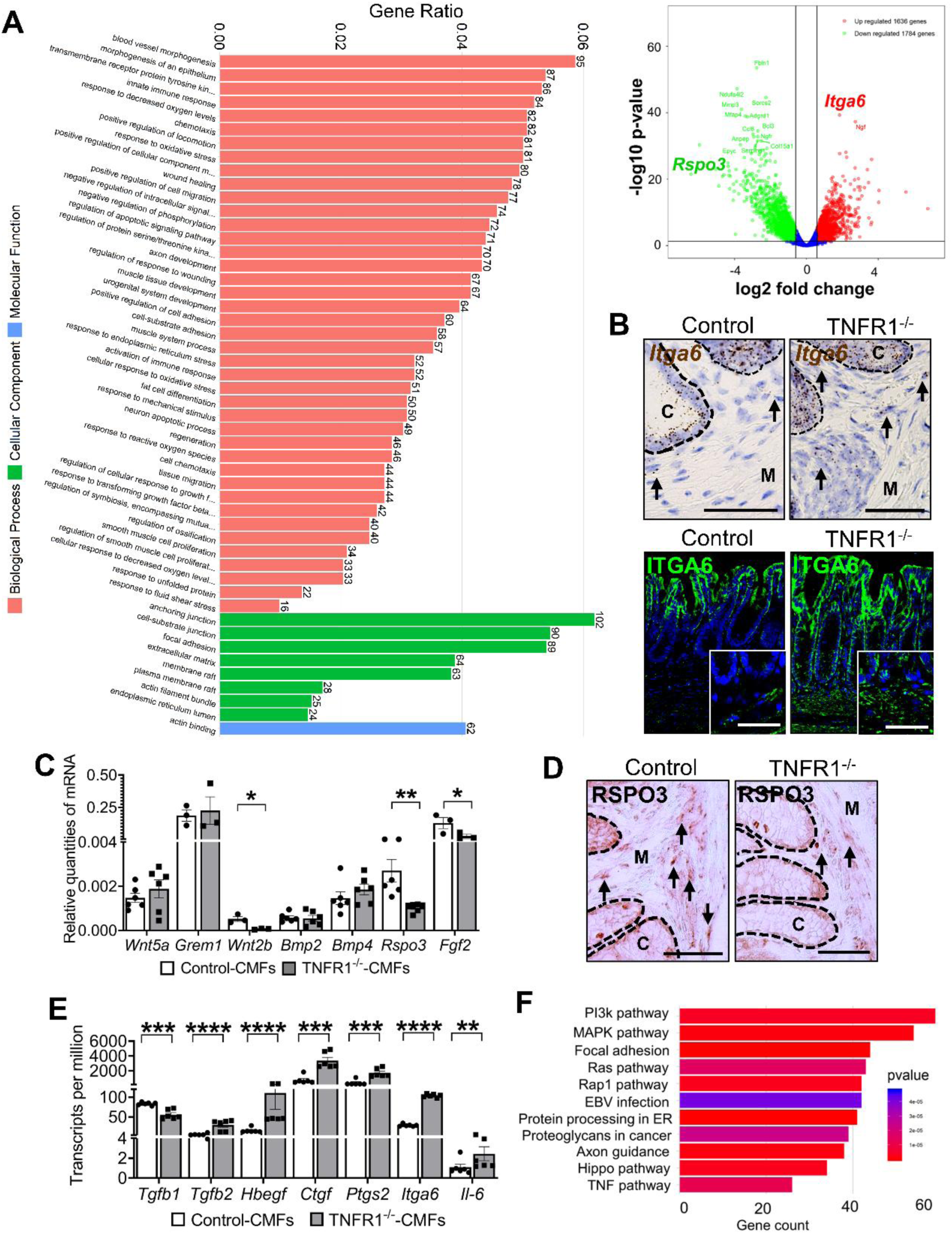
Transcriptome profiling of TNFR1-deficient CMFs identifies upregulation of integrin A6 but loss of the epithelial stem cell niche factor R-spondin 3. (A) KEGG pathway analysis from RNA-Seq showing biological pathways, cellular components, and molecular function of differently expressed genes (DEGs) in TNFR1^-/-^ versus control CMFs, as gene ratios. A volcano plot demonstrating significantly modified DEGs. (B) Localization of ITGA6 in the distal colon of 16-wk-old control and TNFR1^-/-^ mice by *in-situ* hybridization and immunofluorescence staining. Subsets show the localization of ITGA6 in the pericryptal niche cells indicated by black arrows, where C-crypts and M-mesenchyme. (C) q-PCR analysis of Wnt and BMP ligands expressed in control and TNFR1^-/-^ CMFs, at baseline. (D) Localization of RSPO3 in the distal colon of control and TNFR1^-/-^ mice. (E) Genes associated with wound healing and mucosal repair as determined by RNA-Seq of TNFR1^-/-^ versus control CMFs, expressed in transcripts per million. (F) Gene ontology annotation of the top pathways with DEGs sorted by P-value of TNFR1^-/-^ CMFs vs control CMFs. Scale bars = 50µm. *N* = 3 – 6 animals. Error bars are presented as mean ± SEM. Data are analyzed by an unpaired Student t test or a one-way Anova, where * P < 0.05; ** P < 0.01; *** P < 0.001 and **** P < 0.0001.

In contrast to the basal crypt stem cell niche factors, no major differences were observed in the transcript levels of “upper crypt” (i.e., differentiated) niche factors^24^ such as *Wnt5a*, *Bmp2* and *Bmp4* (Figure 3*C*). Several transcripts or pathways associated with wound healing, including *Tgfb2*, *Hbegf*, *Ctgf*, *Ptgs2*, *Itga6,* and *Il6,* and with proliferation and differentiation (e.g., PI3K, MAPK, RAP1, TGF-β, RAS) were upregulated in TNFR1^-/-^ CMFs (Figures 3*E* and 3*F*). We observed downregulation of genes associated with immunity to microbial infections in TNFR1^-/-^ CMFs (Supplementary Figure 5); these genes include *Mmp3*, *Cebpb*, *Icam1*, *Socs3*. Moreover, we observed upregulation of *Jag1* and several other genes (e.g., *Ccl2*, *Csf1r*, *Irf1* and *Irf7*) involved in mucosal repair and in adaptive immune response (Supplementary Figure 5). Thus, deletion of TNFR1 from CMFs alters mesenchymal cell-matrix interactions while downregulating stem cell-associated niche transcript expression and mucosal immunity.

Because the deletion of TNFR1 could theoretically increase compensatory signaling by TNFR2 we assessed for TNFR2 signaling *in vitro* and *in vivo*. The analysis of TNFR1^-/-^ cells failed to recover a signal of TNFR2 enrichment. The levels of TNFR2 transcript and its specific downstream target, Gbp2b^13^ were unchanged in TNF-treated TNFR1^-/-^ CMFs and in TNFR1^-/-^ colon (Supplementary Figures 7A-7C). Furthermore, retrospective analysis showed that the levels of key target genes of mesenchymal TNFR1 – *Tnfr1*, *Mmp9*, *Cd200*, *Rspo3*, *Itga6*, *Pdgfra*, *Acta2*, *Vim*, *Lgr5* and *Ephb3* – were unaltered in TNFR2^-/-^ animals relative to wildtype controls (Supplementary Figure 7D). ^13^ Therefore, the effects of TNFR1 ablation are unlikely to be mediated by compensatory activity of TNFR2.

To assess the relevance of the altered murine TNFR1^-/-^ mesenchyme expression profile to the human UC mesenchyme, we compared gene expression patterns to the mesenchymal cell clusters reported in human UC^15^. Genes enriched in the absence of TNFR1^-/-^ were well represented across all human mesenchymal cell clusters (Supplementary Figures 6A-6C). Formal analysis using GSEA demonstrated that mesenchymal TNFR1 deletion in mice primarily upregulated human myofibroblast (MF) markers while downregulating markers of the UC-associated “S4” mesenchymal cell cluster (Supplementary Figures 6D and 6E). Expansion of the S4 cluster in the human UC mesenchyme was associated with an increase in genes associated with response to TNF, bacterium, and leukocyte migration; ^15^ this resembles the characteristic pathways of “cluster 6” that are incidentally lost in the TNFR1^-/-^ mouse (Supplementary Figure 6D). Although our analysis does not permit establishment of direct equivalency between murine and human cell clusters, the connection between the murine “cluster 6” profile and the human S4 profile seems reasonable given that both exhibit hallmarks of response to TNF/cytokines and might be commonly regulated by TNFR1 signaling. These results support cytokine-driven remodeling of colonic mesenchymal architecture in both mice and humans.

### Inhibiting ITGA6 rescues functional defects in proliferation, wound healing, and stem cell niche factor expression in TNFR1^-/-^ CMFs

We next sought to assess the functionality of PI3K, MAPK and RAP-1 pathways, identified in our transcriptome profiling analysis, for effects on proliferation and wound healing in primary cell cultures. Western blot analyses demonstrated baseline activation of PI3K and MAPK pathways in untreated TNFR1^-/-^ CMFs compared to untreated controls. Levels of phosphorylated AKT and ERK demonstrated a 4-fold increase (Figure 4*A*). The abundance of Ras-related protein 1-GTPase-activating-protein (RAP1-Gap) protein (Figure 4*A*) elevated ∼2-fold in TNFR1^-/-^ CMFs. TNFR1^-/-^ CMFs proliferate at a higher rate (∼20% increase) than controls (Figure 4*B*). Interestingly, there were functional differences between subpopulations of CMFs, with PDGFRα+ cells showing a 23% reduction in proliferation (Figure 4*C*), consistent with their reduced census in TNFR1^-/-^ cultures (cf. Figure 1*C*). In contrast, GLI1 and FOXL1+ cells increased in proliferation in TNFR1^-/-^ CMFs, compared to controls. TNFR1^-/-^ CMFs also exhibited accelerated cell migration, using a 2d scratch wound assay (Figure 4*D*). Therefore, TNFR1 restricts overall mesenchymal proliferation and migration but may selectively promote the PDGFRa+ pericryptal subpopulation.

**Figure 4.**
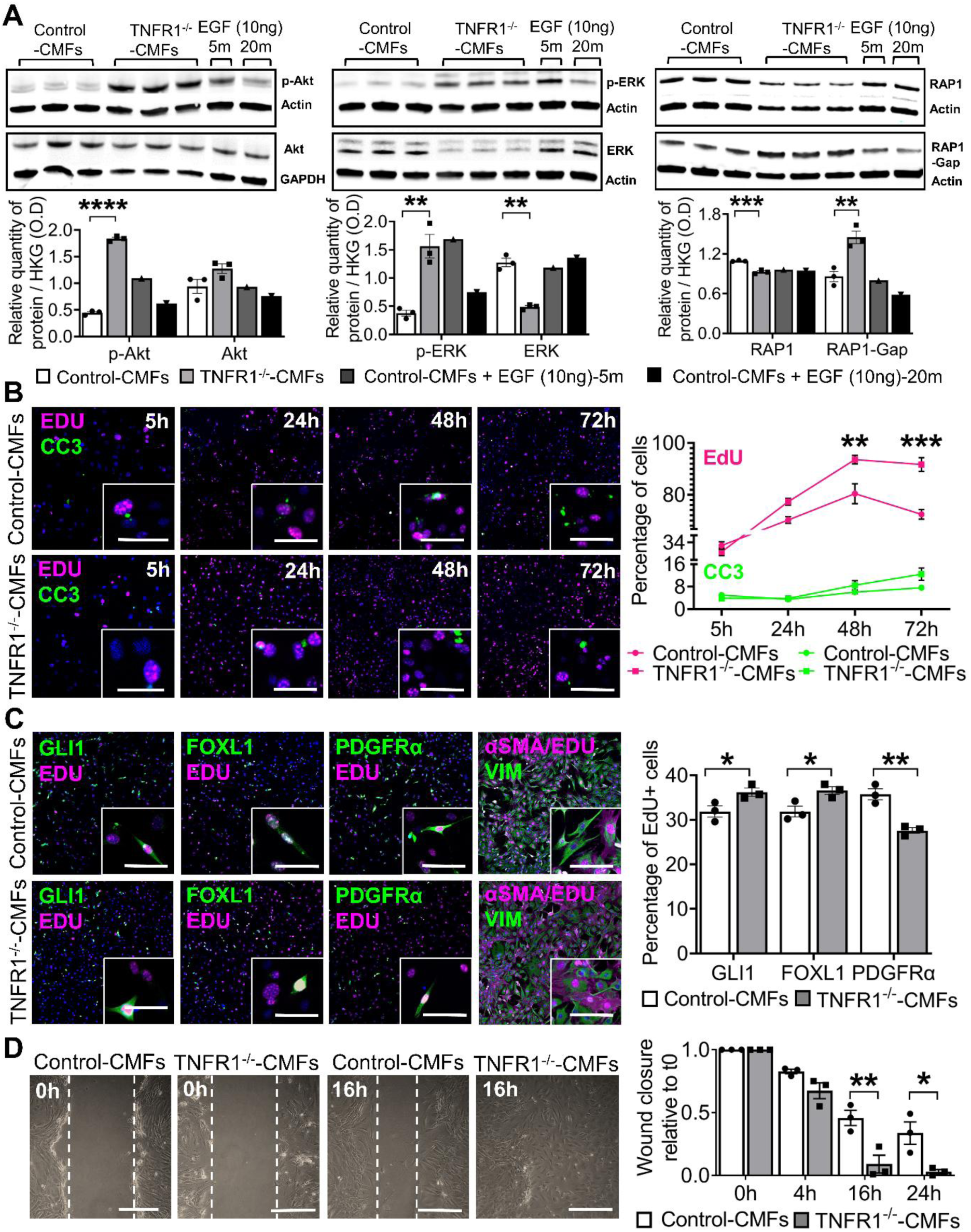
Mesenchymal populations in TNFR1^-/-^ animals show altered proliferation and migration capacities. (A) Activation of the PI3k, MAPK and RAP1 pathways in whole cell lysates of untreated control and TNFR1^-/-^ CMFs, determined by immunoblotting. Positive controls included control CMFs treated with 10 ng/mL EGF for 5- or 20-min. Total protein levels of p-Akt (Ser 473), Akt; p-ERK (Thr202/Tyr204), ERK; and RAP1-Gap, RAP1 was quantified relative to housekeeping genes, Actin or GAPDH. (B) Immuno-co-localization and quantification of the percentage of EdU+ and CC3+ control or TNFR1^-/-^ CMFs over 72 h.(C) Co-localization and quantification of EdU+-FOXL1, PDGFRα, and GLI1+ control or TNFR1^-/-^ CMFs at 72 h. Scale bars = 25µm. (D) Cell migration as determined by wound scratch assays of untreated control and TNFR1^-/-^ CMFs at baseline, over 0 h – 24 h. Migrated cells were quantified by determining the wound closure rate relative to t 0. Scale bars = 50µm. All images are representative of *N* = 3 – 6 animals. Error bars are presented as mean ± SEM. Data are analyzed by an unpaired Student *t* test or a one-way Anova, where * P < 0.05; ** P < 0.01; *** P < 0.001 and **** P < 0.0001.

Next, we defined a role for ITGA6 in the regulation of proliferation and migration of TNFR1^-/-^ CMFs. Treatment of TNFR1^-/-^ CMFs with anti-ITGA6 blocking antibody (Supplementary Figure 8A) restored ERK and RAP1 signaling (Figure 5*A*), cellular proliferation (Figure 5*B*), and wound closure rate (Figure 5*C*) to near-control levels. Since the crosstalk between integrins and PDGFRs is essential to effective growth factor signaling, ^25^ we next assessed ITGA6’s role in stem cell niche function. Anti-ITGA6 antibody treatment of TNFR1^-/-^ CMFs restored the numbers of PDGFRα+ cells (Figure 5*D*), expression of PDGFRα protein, and expression of RSPO3 to near-control levels (Figure 5*E*). Thus, ITGA6 forms a central “hub” of integrin-matrix signaling that mediates TNFR1’s diverse functions within the different mesenchymal populations.

**Figure 5.**
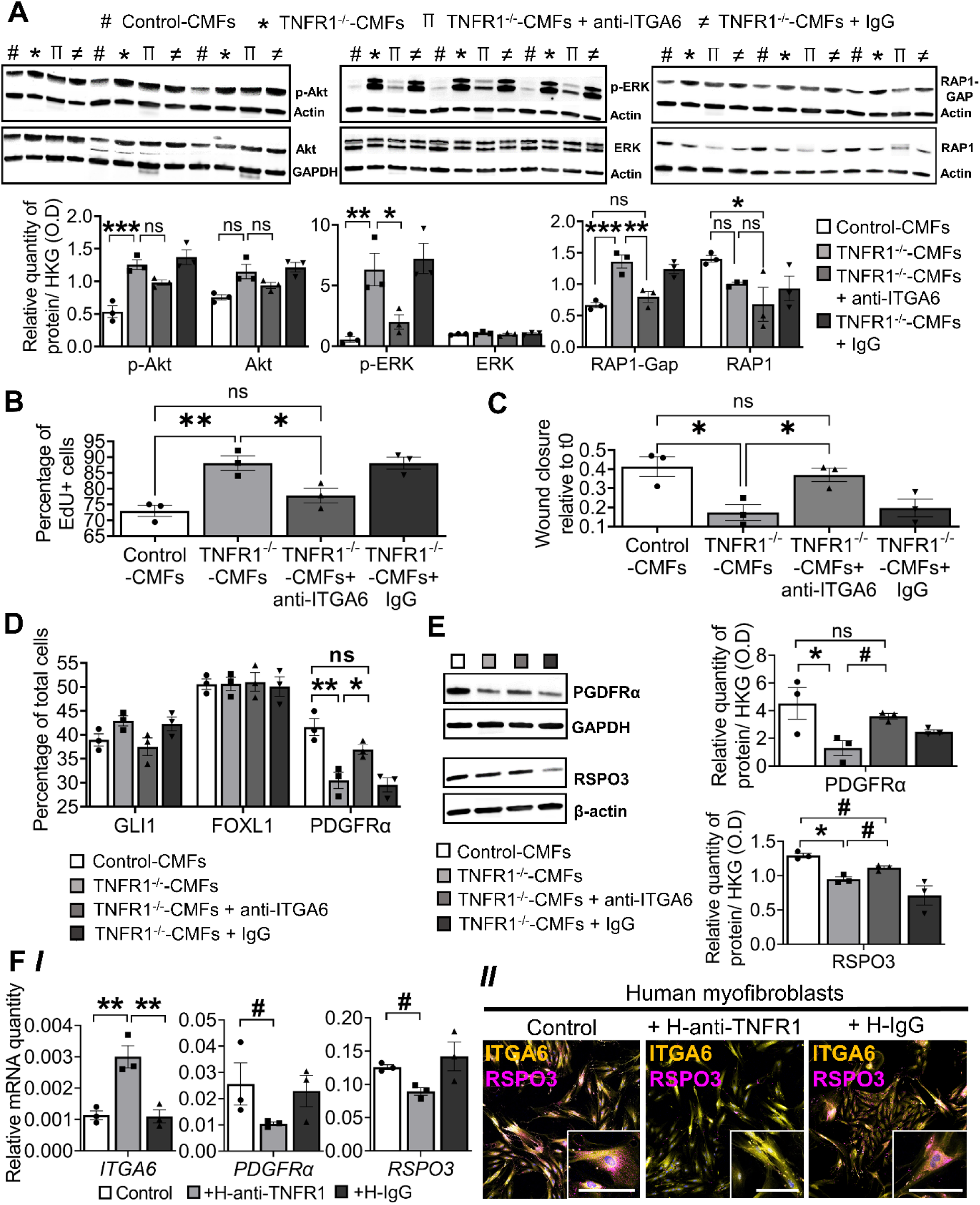
Inhibiting ITGA6 rescues functional defects in proliferation, wound healing, and stem cell niche factor expression in TNFR1^-/-^ CMFs. (A) Activation of the PI3k, MAPK and RAP1 pathways in whole cell lysates of TNFR1^-/-^ CMFs treated with 0.2 µg anti-ITGA6 antibody and IgG control, relative to untreated control CMFs, determined by immunoblotting. The total protein levels of p-Akt, Akt; p-ERK, ERK and RAP1-Gap, RAP1 was quantified relative to housekeeping genes, Actin or GAPDH. *N* = 3 animals. (B) Quantification of the percentage of EdU+ TNFR1^-/-^ CMFs or (C) Cell migration determined by wound scratch assays of TNFR1^-/-^ CMFs treated with 0.2 µg anti-ITGA6 antibody and IgG control, relative to untreated control CMFs, over 0 h – 16 h. (D) Quantification of pericryptal mesenchymal cell markers, FOXL1, PDGFRα, and GLI1 or (E) Quantification of total protein levels of PDGFRα and RSPO3 in TNFR1^-/-^ CMFs treated with 0.2 µg anti-ITGA6 antibody and IgG control, relative to untreated control CMFs. (F) (*I*) q-PCR analysis of *ITGA6*, *PDGFRα* and *RSPO3*, and (*II*) localization of ITGA6 and RSPO3 in primary human colonic myofibroblasts, 18Co, treated with 0.5 µg/mL anti-TNFR1 and its IgG isotype control for 5 d. Scale bars = 25 µm. All images are representative of *N* = 3 animals. Error bars are presented as mean ± SEM. Data are analyzed by a one-way Anova, where * P < 0.05; ** P < 0.01; *** P < 0.001, and ^#^ P = 0.05-0.09.

To determine whether these findings translate to human intestinal mesenchyme, we treated human primary 18Co myofibroblasts with 0.5 µg of anti-TNFR1 antibody (Supplementary Figures 8B-8D). Treated cells exhibited significant increase in the transcript levels of *ITGA6* and a downregulation of *PDGFRA* and *RSPO3* (Figure 5*FI*), similar to mouse TNFR1^-/-^ CMFs. We show that ITGA6 and RSPO3 co-localize in human primary 18Co myofibroblasts and that treatment with the anti-TNFR1 antibody reduces the balance of RSPO3/ITGA6 expression (Figure 5*FII*). Therefore, TNFR1 regulates ITGA6 and the stem cell niche factors, PDGFRα and RSPO3, in human colonic mesenchymal cells.

### RSPO3 expression by mesenchymal cells is required for homeostatic expression of stem cell numbers and function in TNFR1^-/-^ epithelium

Because TNFR1^-/-^ CMFs have reduced expression of niche factors including RSPO3, we tested their functional role by establishing a unique model of epithelium and mesenchyme co-culture. By co-culture of control or TNFR1^-/-^ colonoids with control or TNFR1^-/-^ mesenchymal cells in a 3D matrix (Figure 6*A*), we can study the effect of a specific genotype on epithelial and mesenchymal expression and function. Co-culture of control or TNFR1^-/-^ organoids with TNFR1^-/-^ CMFs significantly reduced total mRNA expression of *Lgr5* and *Ephb3* (an alternate stem cell marker^26, 27^) by 2-fold (P = 0.02), compared to co-culture of control or TNFR1^-/-^ organoids with control CMFs (Figure 6*BI*). We confirmed this observation (i.e., ∼30% reduction of Lgr5+ cells in presence of TNFR1^-/-^ CMFs) by co-culturing control or TNFR1^-/-^ CMFs with colonoids derived from Lgr5::DTR-EGFP mice, which label Lgr5+ stem cells with GFP and permit direct visualization of Lgr5+ cells via confocal microscopy (Supplementary Figures 9A and 9B). Moreover, co-culture with TNFR1^-/-^ CMFs altered the morphology of colonoids, resulting in a 40% decrease in the proportion of budding colonoid structures, with a parallel increase in a rapidly expanding monolayer-like epithelium (Figure 6*BII*). Thus, mesenchymal TNFR1 regulates the epithelial stem cell compartment and functionally alters the growth pattern of colonoids.

**Figure 6.**
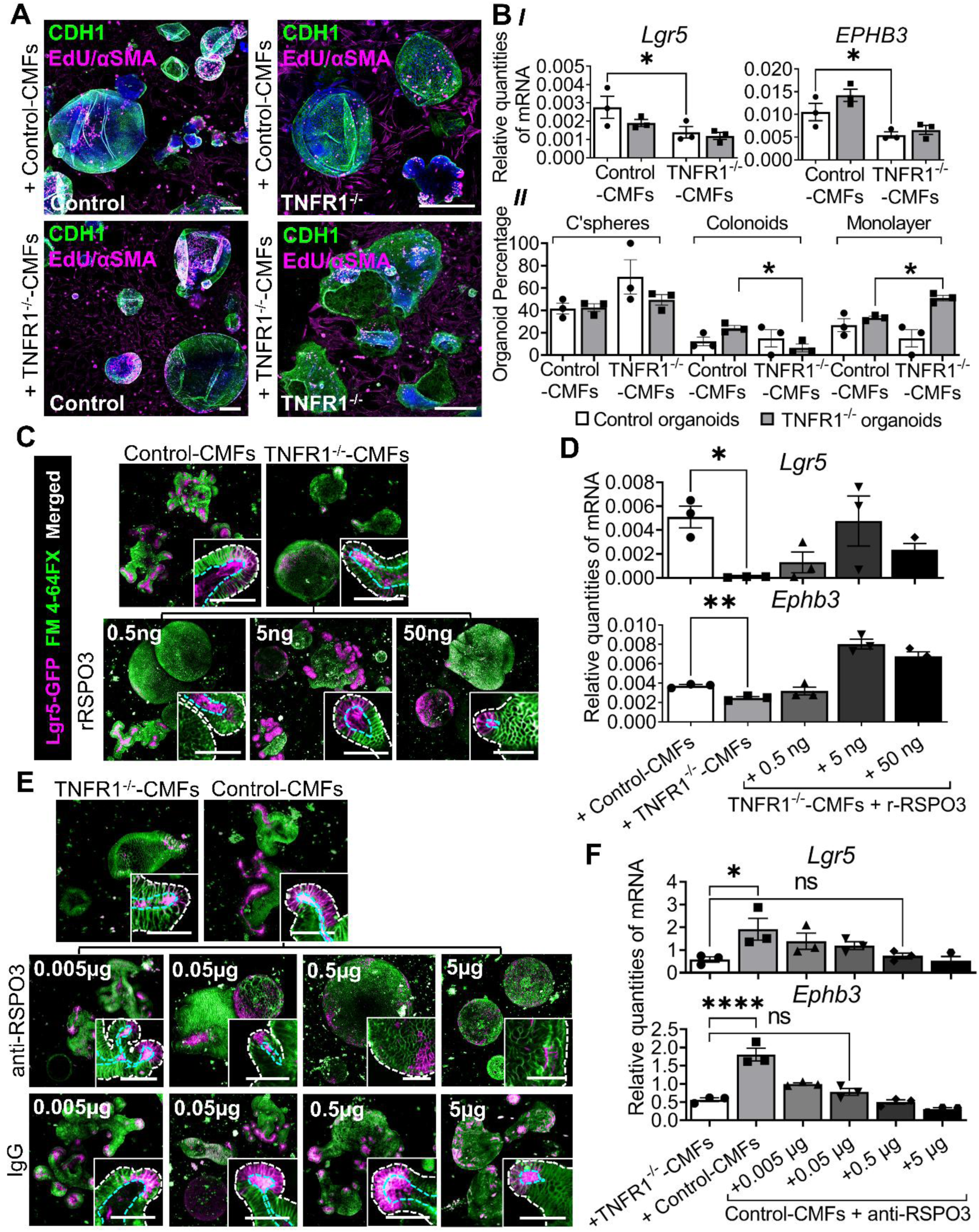
RSPO3 expression by mesenchymal cells is required for homeostatic expression of stem cell numbers and function in TNFR1^-/-^ epithelium. (A) Immunofluorescence staining of epithelial E-cadherin (CDH1); EdU+ and aSMA+ CMFs in co-cultures of control or TNFR1^-/-^ organoids with control or TNFR1^-/-^ CMFs. (B) qPCR analysis of (*I*) *Lgr5* and *Ephb3* mRNA expression, and (*II*) quantification of colonospheres, colonoids and colonoid monolayers, in co-cultures of control or TNFR1^-/-^ organoids with control or TNFR1^-/-^ CMFs. (C) Immunofluorescence staining of GFP and FM 4-64X, and (D) q-PCR analysis of *Lgr5* and *Ephb3* levels in Lgr5-GFP organoids co-cultured with untreated control or TNFR1^-/-^ CMFs and TNFR1^-/-^ CMFs treated with increasing concentrations of recombinant-RSPO3 (0.5 - 50 ng/mL) for 5 d in culture. (E) Immunofluorescence staining of GFP and FM 4-64X, and (F) q-PCR analysis of *Lgr5* and *Ephb3* levels in Lgr5-GFP organoids co-cultured with untreated TNFR1^-/-^ or controls CMFs and controls CMFs treated with increasing concentrations of anti-RSPO3 antibody and its IgG isotype control (0.005 - 5 µg/mL) for 5 d in culture. The cyan dotted line represents the cell apical border, and the white line denotes the basal border. Scale bars = 50 µm. All images are representative of *N* = 3 animals. Error bars are presented as mean ± SEM. Data are analyzed by a one-way Anova, where * P < 0.05; ** P < 0.01; and **** P < 0.0001.

We next asked whether RSPO3 was sufficient to restore functional support of Lgr5+ epithelial cells in the presence of TNFR1^-/-^ CMFs. Colonoids from Lgr5::DTR-EGFP mice were co-cultured with TNFR1^-/-^ CMFs supplemented with increasing concentrations of recombinant RSPO3 for 5 days. Treatment with recombinant RSPO3 at a dose of 5 ng increased the number of crypt-like budding structures in colonoids (Figure 6*C*) and restored expression to *Lgr5* and *Ephb3* to control levels (Figure 6*D*). However, treatment with higher RSPO3 concentrations (50 ng) resulted in a loss of crypt like budding structures consistent with an undifferentiated epithelial phenotype. Concurrently, the levels of *Lgr5* and *Ephb3* were also reduced in these colonoids. To confirm the specificity of recombinant RSPO3 in restoring stem cell marker expression, control CMFs were treated with an anti-RSPO3 antibody, and stem cell expression in organoid co-cultures were measured. Anti-RSPO3 inhibited *Lgr5* and *Ephb3* expression in a dose-dependent manner, such that at the 0.05 µg dose, the levels of *Lgr5* and *Ephb3* expression are comparable to those found in co-cultures with TNFR1^-/-^CMFs (Figures 6*E* and 6*F*). Therefore, RSPO3 rescues the loss of TNFR1 signaling in mesenchymal cells with regard to epithelial stem cell marker expression.

### TNFR1-deficient mice have reduced Lgr5 and Ephb3 stem cell marker expression

To determine if TNFR1 expression is required for basal stem-cell marker expression *in vivo*, we compared the colons of TNFR1^-/-^ mice to their heterozygote littermate controls (TNFR1^+/-^). Consistent with our *in vitro* findings, there was a 44% reduction in Lgr5+ stem numbers and a ∼2-fold decrease in Lgr5 mRNA expression (Figure 7*AI*) in TNFR1^-/-^ animals. Similar results were obtained from analysis of *Ephb3* expression (Figure 7*AII*).

**Figure 7.**
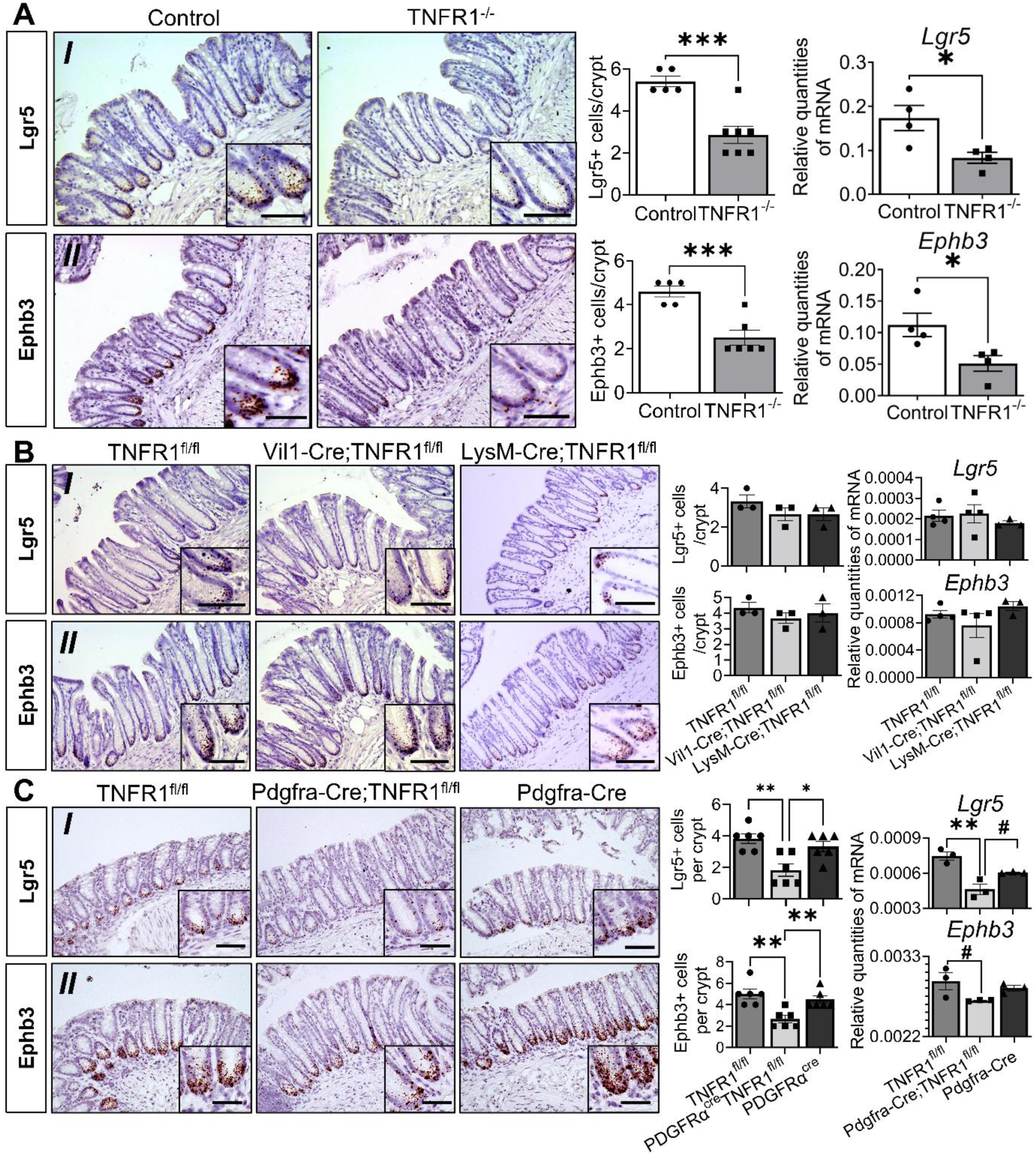
TNFR1 deletion in colonic pericryptal PDGFRα+ cells but not intestinal epithelial or myeloid cells results in a loss in stem cell marker expression. (A) Localization and quantification of (*I*) Lgr5+ cells per crypt and qPCR analysis of *Lgr5* expression, or (*II*) Ephb3+ cells per crypt and qPCR analysis of *Ephb3* expression from the distal colon of control (TNFR1^+/-^) and TNFR1^-/-^ mice. (B) Localization and quantification of (*I*) Lgr5+ cells per crypt and qPCR analysis of *Lgr5* expression, or (*II*) Ephb3+ cells per crypt and qPCR analysis of *Ephb3* expression from the distal colon of TNFR1^fl/fl^, Vil1-Cre;TNFR1^fl/fl^, and LysM-Cre;TNFR1^fl/fl^ mice. (C) Localization and quantification of (*I*) Lgr5+ cells per crypt and qPCR analysis of *Lgr5* expression, or (*II*) Ephb3+ cells per crypt and qPCR analysis of *Ephb3* expression from the distal colon of TNFR1^fl/fl^, Pdgfra-Cre;TNFR1^fl/fl^, and Pdgfra-Cre mice. Subsets show the punctate staining of Lgr5 and Ephb3 in the crypt base cells. Scale bars = 50 µm. All images are representative of *N* = 3 – 6 animals. Error bars are presented as mean ± SEM. Data are analyzed by an unpaired Student *t* test, or a one-way Anova, where * P < 0.05; ** P < 0.01; *** P < 0.001; and ^#^ P = 0.05-0.09.

To determine if the effects *in vivo* could be explained by tissue-specific function of TNFR1, we compared expression in mice with an intestinal epithelial specific deletion of TNFR1 (Vil1-Cre;TNFR1^fl/fl^), myeloid cell specific deletion of TNFR1 (LysM-Cre;TNFR1^fl/fl^), or PDGFRα+ cell specific deletion of TNFR1 (Pdgfra-Cre;TNFR1^fl/fl^). No significant differences in numbers of Lgr5 (Figure 7*BI*) or Ephb3+ stem cells (Figure 7*BII*) were observed between Vil1-Cre;TNFR1^fl/fl^, LysM-Cre;TNFR1^fl/fl^, and their TNFR1^fl/fl^ controls. However, Pdgfra-Cre;TNFR1^fl/fl^ animals demonstrated a significant decrease in Lgr5+ (Figure 7*CI*) and Ephb3+ (Figure 7*CII*) stem cells. Thus, the effects of global TNFR1 deletion on the epithelial stem cell pool *in vivo* are not explained by epithelial or myeloid cell-autonomous effects, but by the specific regulation of the pericryptal PDGFRα+ mesenchyme. Hence, our findings demonstrate a role for mesenchymal TNFR1 expression in regulating the intestinal epithelial stem cell pool via epithelial-pericryptal mesenchymal cell interactions.

## Discussion

The profusion of therapies that inhibit TNF signaling in the context of autoimmunity necessitates a fuller understanding of their potential physiological impact. This is a critical need due to the high documented rates of clinical non-response in conditions such as IBD. ^4^ Studies have demonstrated “beneficial” roles of TNF in IBD^8–11^ and reported new-onset IBD with use of anti-TNF agents, ^28^ consistent with a possible anti-inflammatory role for TNF in some contexts. Although previous studies have supported a role for TNF signaling in maintenance of epithelium and healing responses in murine intestinal inflammation models, the mechanisms have remained unknown and have largely been assumed to be epithelial cell-autonomous. ^8–11^ Our study shows that TNFR1-mediated signaling is central to the proper specification of colonic pericryptal mesenchymal cells and to the fundamental mesenchymal-epithelial paracrine axis that maintains the crypt stem cell pool. This cytokine signal is essential in shaping mesenchymal cellular diversity of the colon. Combined with recent work showing TNFR1’s role in mediating mucosal tolerance to the microbiome in preclinical IBD models, ^11^ our study outlines a multimodal mechanism, with mesenchymal cells regulating the epithelial response to cytokines, through which TNFR1 signaling might enforce mucosal homeostasis. In this study, the genetic or antibody-mediated depletion of TNFR1 signaling had profound effects on mesenchymal cell function and epithelial stem cell census, key aspects of mucosal health. Future studies could explore how these intercellular networks are altered in patients treated with anti-TNF agents.

We report that mesenchymal TNFR1 is required to preserve the overall diversity of colonic mesenchymal populations at homeostasis, including a sub-population of pericryptal cells that are uniquely enriched for TNF and IFN signaling genes. TNFR1 ablation *in vivo* alters the proportions of mesenchymal cell subpopulations (Figure 1). As there are several known markers of mesenchymal subpopulations which potentially overlap, we performed single-cell RNA transcriptomics to precisely enumerate the diversity of colonic mesenchymal cell-types. Moreover, as the “cytokine storm” of IBD is associated with broad changes in mesenchymal cell differentiation^15, 19–21^ we had anticipated that synergy between many cytokines could drive these changes. However, in single-cell experiments, in Figure 2, we found that the deletion of a single cytokine receptor results in fundamental changes such that TNFR1^-/-^ and TNFR1^+/-^ cell clusters were mostly mutually exclusive. Although cluster exclusion can sometimes arise from batch effects, these analyses were performed as a single batch with independent cultures from co-housed littermate mice. Thus, we believe the results are faithful to the composition of the cultures; however, we cannot rule out elimination or modification, due to culturing, of certain cell types initially present *in vivo*. However, our analysis, on several occasions, demonstrates that *in vitro* cultures faithfully recapitulated the alterations in the major mesenchymal phenotypes observed *in vivo*, and this was further validated by both single-cell and bulk RNA-seq data. Multiple approaches confirmed that a mesenchymal cell-type appears to be downregulated in TNFR1^-/-^ mice. This cell-type is located near the base of the crypt, expresses Mmp9 and Cd200 markers, and has a transcriptional profile with hallmarks of TNF signaling. The cell-type appears to be a subset of PDGFRα+ cells and shares some similarity with a UC-associated mesenchymal cell type. ^15^ Future studies will need to characterize how this cell-type integrates with other CMF subpopulations in terms of hierarchy and niche function, as well as determine whether it has specific functions that distinguish it from other PDGFRα+ cells. Taken together, these results demonstrate that TNF is an essential driver of CMF specification and function.

Our study defines a novel pathway of TNFR1-driven repression of integrins (ITGA6), the result of which is differential alteration of cell proliferation and wound healing-associated pathways within the mesenchymal subpopulations (Figures 3-5). Moreover, we found that the TNFR1^-/-^ CMFs upregulating ITGA6 fail to perform one of their key functions: the secretion of sufficient niche-specific growth factors to sustain the epithelial stem cell pool^25^ (Figures 5 and 6). TNFR1’s effects on the stem cell niche converge to be mediated by RSPO3. Supplementation of TNFR1^-/-^ CMFs with exogenous RSPO3 *in vitro* restored the number of Lgr5+ stem cells to control levels (Figure 6). RSPO3, the most highly expressed intestinal Wnt ligand, is crucial for sustaining the Lgr5+ stem cell pool and is known to be entirely produced by pericryptal niche cells. ^22, 23, 29^ Consistent with our findings, PDGFRα+ pericryptal mesenchymal cells have been identified as a key pericryptal source of RSPO3^23^ and support epithelial architecture and the renewal of stem cells during homeostasis and the injury response. ^30, 31^ Hence, the development of approaches to restore RSPO3-mediated signaling or inhibit ITGA6 may partially compensate for the effects of TNFR1 ablation on mesenchymal-epithelial interactions and the stem cell niche.

Deletion of mesenchymal TNFR1 has profound effects on epithelial stem cells *in vivo* and *in vitro*. We show in Figure 7 that animals with global and mesenchymal-specific TNFR1 depletion exhibit a loss of Lgr5+ stem cells. That this occurs without attendant changes in crypt height suggests the presence of other homeostatic mechanisms to maintain the overall crypt surface area. The indirect mechanism through which mesenchymal TNFR1 regulates the epithelium through the niche reinforces the need for cell culture systems that partially reconstitute heterotypic cellular interactions. We believe the epithelium and mesenchyme co-culture model demonstrated here is a good example of this. Co-culture systems will become increasingly important for investigation of cytokine signaling pathways, as these components are expressed by multiple tissue types in the mucosa, and for resolution of the mechanistic basis of inflammatory diseases.

## Materials and Methods

Additional details of experimental procedures and reagents are available as a supplement to this manuscript.

### Experimental Animals

Animals were cared for according to CHLA IACUC rules, protocol #288. All animals were purchased from Jackson Laboratory, except for Lgr5::DTR-EGFP (gift from Fred de Sauvage, Genentech, USA) and TNFR1^fl/fl^ (gift from George Kollias, Alexander Fleming Biomedical Sciences Research Center, Vari, Greece). Experimental and control mice were co-housed for > 4 weeks prior to experimentation. Both females and males (6-16 wks-old) were used.

### Murine Three-Dimensional Organoid and Myofibroblasts Co-culture

Distal colonic tissue isolated from control, TNFR1^-/-^ or Lgr5::DTR-EGFP mice were processed for digestion and crypt isolation and cultured in Matrigel and Intesticult (Stem cell Technologies), for organoid generation as per previously established protocols. ^11, 32^ Passaged organoids were broken into single cells via syringe extrusion. Epithelial cells were mixed with CMFs at a ratio of 10:1 into Matrigel. Co-cultures were grown and maintained in 60% IntestiCult™ (STEMCELL, Seattle, WA; no. 06005) and 40% CMFs-growth medium at 37°C. Cultures were treated with recombinant RSPO3 (R&D systems, Minneapolis, MN; no. 4120-RS), an anti-RSPO3 antibody (R&D systems, no. AF35001), an IgG isotype control (R&D systems, no. 5-001-A), or anti-ITGA6 antibody (Sigma, St. Louis, MO; no. MAB1378) and IgG isotype control (Millipore Sigma, no. 12-370). Organoids were stained with FM4-64FX membrane dye prior to confocal imaging.

### RNA-Seq

mRNA transcripts from control and TNFR1^-/-^ CMFs were analyzed from RNA extracts via 2x75 bp sequencing on a HiSeq 4000 (Illumina). Samples include 6 control and 6 TNFR1^-/-^ primary CMF cultures (passage numbers 6-8) that were originally isolated from distinct mice and amplified *in vitro* in Matrigel. Transcript abundance was calculated using RSEM v1.2.15. Differentially expressed genes were identified using the edgeR program. Genes showing altered expression with p < 0.05 and more than 1.5-fold changes were considered differentially expressed.

### scRNA-Seq

Mesenchymal cells were analyzed using the standard Chromium (10x Genomics) pipeline with a targeted recovery of 10,000 cells per sample. A total of 6 samples (3 control and 3 TNFR1^-/-^) representing individual CMF cultures from distinct mice were isolated and amplified in vitro in Matrigel (passage number > 3). Cells were dissociated into a single-cell suspension and pooled libraries were sequenced on an Illumina HiSeq 4000 machine to obtain an estimated read depth of 50,000 reads/cell. The resulting reads were analyzed in cellranger 5.0.0 using the mm10-3.0.0 reference genome. Matrix abundance files were merged and loaded into monocle3 for downstream analysis. Cells with <100 UMI counts were filtered out.

### Statistical Analysis

GraphPad Prism software was used for statistical analyses. Data are presented as means with standard error of mean (SEM). Analysis was carried out by either an unpaired Student *t* test or a one-way Anova, with a Šídáks post-hoc correction, as indicated in figure legends, where * P < 0.05; ** P < 0.01; *** P < 0.001; **** P < 0.0001; and ^#^ P=0.05-0.09.

## Acknowledgements

This study was supported by the U.S. National Institutes of Health (RO1-DK056008 and RO1-DK108648, to D. Brent Polk) and by the San Diego Digestive Diseases Research Center (P30-DK120515). CYL is supported by a career development award from the Crohn’s and Colitis Foundation (#694110) and by the University of Chicago, Digestive Diseases Research Core Center (P30-DK042086). The library preparation and sequencing for the single cell- and RNA Sequencing data were performed with the resources and expertise of Children’s Hospital Los Angeles Single Cell, Sequencing, and CyTOF Core Laboratory (Los Angeles, CA).

## Transcript Profiling

The single cell RNA Sequencing dataset generated during this study is available at NCBI Gene Expression Omnibus with the accession number GSE180346. Full analysis scripts are available at https://github.com/stalepig/2021-mesenchyme-scrnaseq. The RNA Sequencing dataset for control and TNFR1^-/-^ CMFs (colonic myofibroblasts) is available at NCBI Gene Expression Omnibus, identified by the accession number GSE180421.

## Data Transparency

Data, analytic methods, and study materials will be made available from the corresponding author on reasonable request.

## Abbreviations

TNFR: Tumor Necrosis Factor Receptor
TNF: Tumor Necrosis Factor
IBD: Inflammatory Bowel Disease
MSCs: Mesenchymal Cells
RNA-Seq: RNA Sequencing
scRNA-Seq: single cell RNA Sequencing
CMFs: Colonic Myofibroblasts
C’spheres: Colonospheres
EdU: 5-ethynyl-2’-deoxyuridine
CC3: Cleaved Caspase 3
No: Number
^fl/fl^: floxed/floxed
Vil1-Cre;TNFR1^fl/fl^: Deletion of TNFR1 in intestinal epithelial cells
LysM: Cre;TNFR1^fl/fl^ Deletion of TNFR1 in myeloid cells
Pdgfra-Cre;TNFR1^fl/fl^: Deletion of TNFR1 in PDGFRα+ cells
αSMA: alpha smooth muscle actin
GLI: GLI family zinc finger 1
FOXL1: forkhead box L1
FM 4-64FX: N-(3-Triethylammoniumpropyl)-4-(6-(4-(Diethylamino) Phenyl) Hexatrienyl) Pyridinium Dibromide
VIM: Vimentin
PDGFRα: Platelet derived growth factor receptor alpha
ERK: Extracellular-signal-regulated kinase
RAP1: Ras-proximate-1 or Ras-related protein 1RAP1-GAP Ras-proximate-1 or Ras-related protein 1-GTPase-activating-protein
ITGA6: Integrin A6
RSPO3: R-spondin 3
IgG: Immunoglobulin G
r-RSPO3: recombinant-R-spondin 3
P13k: Phosphoinositide 3-kinase
MAPK: Mitogen-activated protein kinase
TGF-β: Transforming growth factor beta
DEGs: Differently expressed genes.

## Author Contributions

Conceptualization, S.G., C.Y.L., and D.B.P.; Methodology, S.G., C.Y.L., N.G., Y.H., and D.B.P.; Software, C.Y.L.; Validation, S.G., C.Y.L., N.G., Y.H., and D.B.P.; Formal Analysis, S.G., and C.Y.L.; Investigation, S.G., C.Y.L., and N.G.; Resources, S.G., C.Y.L., N.G., Y.H., K.M.M., T.C.G., and D.B.P.; Data Curation, S.G., C.Y.L., N.G., and D.B.P.; Writing – Original Draft, S.G.; Writing – Review & Editing, S.G., C.Y.L., and D.B.P.; Visualization, S.G., C.Y.L., and D.B.P.; Supervision, S.G., C.Y.L., and D.B.P.; Project Administration, D.B.P.; Funding Acquisition, C.Y.L., and D.B.P.

## Synopsis

TNF-TNFR1 signaling on pericryptal platelet-derived growth factor receptor alpha (PDGFRα)+ mesenchymal cells maintains colonic Lgr5+ epithelial stem cells, the overall colonic mesenchymal cell diversity, and a distinct sub-population of pericryptal mesenchymal cells enriched for TNF and IFN signaling genes.

## Supplementary Methods and Materials

### Experimental Animals

Animals were cared for as per Institutional Animal Care and Use Committee (IACUC) rules and regulations at Children’s Hospital Los Angeles (CHLA) under internal protocol number 288. Preceding dissection, animals were humanely anesthetized with isoflurane and euthanized by cervical dislocation. C57BL/6J-WT (cat# 000664), TNFR1^-/-^ (cat# 002818), Pdgfra-Cre (cat# 013148), LysM-Cre (cat# 004781), and TNFR2^-/-^ (cat# 002620) mice were obtained from Jackson Laboratory (Maine, ME). Lgr5::EGFP-IRES-CreERT2 (Fred de Sauvage, Genentech, USA), and TNFR1^fl/fl^ (George Kollias, Alexander Fleming Biomedical Sciences Research Center, Vari, Greece) mice were kindly gifted. Mesenchymal Pdgfra-Cre mice were crossed to TNFR1^fl/fl^ mice to generate Pdgfra-Cre;TNFR1^fl/fl^ animals. TNFR1 deletion in mesenchymal PDGFRα+ cells were confirmed by genotyping (Supplementary Figure 1AI) and the loss of total protein levels of TNFR1 (Supplementary Figure 1AII) in the primary mesenchyme but not the organoids derived from 3 independent Pdgfra-Cre;TNFR1^fl/fl^, TNFR1^fl/fl^, and Pdgfra-Cre animals. Intestinal epithelial-specific TNFR1 knockout mice (Vil1-Cre;TNFR1^fl/fl^) were generated by crossing Villin-CreER mice^33^ with TNFR1^fl/fl^ mice, and myeloid cells-specific TNFR1 knockout mice (LysM-Cre;TNFR1^fl/fl^) were generated by crossing LysM-Cre with TNFR1^fl/fl^ mice. To maintain proper experimental controls, littermate controls of transgenic mice were co-housed for > 4 weeks prior to experimental collection and analysis. Adult male and female mice (6/12-16 weeks) were involved in the study.

### Mice Tissue Collection

Colon and cecum were dissected out from euthanized mice and cleared of fecal contents. For histology, the colon was swiss-rolled and fixed in 10% neutral buffered formalin. 5 μm slices of the tissue were sectioned. The distal end of the colon (2cm from the anus) was longitudinally split into half and processed for RNA and protein isolation and in the same manner for generation of organoids and primary mesenchymal cells.

### Murine Three-Dimensional Organoid Culture

Distal colonic tissue isolated from control (TNFR1^+/-^), TNFR1^-/-^ or Lgr5::EGFP-IRES-CreERT2 mice were processed for digestion and crypt isolation as per previously established protocols. ^11, 32^ For passage, organoids were broken mechanically by syringing using a 27 G X 1/2 needle (Becton Dickinson, San Diego, CA, cat# 305109), then transferred to fresh Matrigel. The passage was performed every 5–8 days with a 1:4 split ratio and medium was changed after every 2-3 days. Each experiment was repeated thrice. Each condition was examined in triplicate with multiple (>30-50) organoids in each sample.

### Co-culture Experiments

#### Isolation of Primary Colonic Myofibroblasts

Primary mouse colonic myofibroblasts (CMFs) were isolated and maintained as described previously. ^16^ Briefly, primary colonic myofibroblasts (CMFs) isolated from the distal end of 16-wk-old control (TNFR1^+/-^) and TNFR1^-/-^ mice were repeatedly stirred at 250 rpm in 5 mM EDTA/HBSS medium (without Ca^++^ and Mg^++^) at 37°C for 15 min for a total of 5 washes. Following two washes of ice-cold PBS, the tissue was digested in DMEM, low glucose, GlutaMAX™ Supplement, pyruvate (Gibco, Waltham, MA; cat# 10567014) plus Penicillin-Streptomycin (10,000 U/mL) with, 0.31 mg/mL Dispase (Gibco, cat# 17105041) and 0.25 mg/mL collagenase Type XI (Gibco, cat# 17104019) for up to 60 minutes at 250 rpm at 37°C, until the tissue appeared stringy. Digested tissue was treated with ACK lysis buffer (4.15 g NH_4_Cl, 0.5 g KHCO_3_, 18.6 mg Na_2_EDTA, 400 ml H_2_O, pH 7.2-7.4) for 5 min and then passed through a 100 μm cell strainer into 6-well dishes of CMF-growth medium. Primary cultures were maintained in a CMF-growth medium comprised of DMEM, low glucose, GlutaMAX™ Supplement, pyruvate (Gibco, cat# 10567014) plus 10,000 U/mL Penicillin-Streptomycin (cat# 15140163) with 10% FBS (Corning, Corning, NY; cat# 35-010-CV) and 1X recombinant human insulin, human transferrin and sodium selenite (ITS) solution (Sigma, St. Louis, MO; cat# I3146), and 20 ng/mL recombinant murine epidermal growth factor (Peprotech, Rocky Hill, NJ; cat# 315-09) at 37°C in a 5% CO_2_/air mix. After 24 h incubation, non-adherent cells were washed away. CMFs were grown in two-dimensional culture for amplification of cells for the main experimental setup. Control or TNFR1^-/-^ primary CMFs were seeded at a density of 1x10^3^ cells/mL in parallel in 6-well plates and were grown to 70% confluence. At day 5, CMFs in two-dimensional culture recovered similar numbers of mesenchymal populations (VIM+, αSMA+, PDGFRα+, GLI1+, or FOXL1+) as CMFs in three-dimensional culture at day 5 (Supplementary Figure 4B).

#### TNF treatment of primary colonic myofibroblasts

Control or TNFR1^-/-^ primary CMFs were seeded at a density of 1x10^3^ cells/mL in parallel in 6-well plates and were grown to 70% confluence. Cells were starved for 24 h in DMEM, low glucose, GlutaMAX™ Supplement, pyruvate medium plus 10,000 U/mL Penicillin-Streptomycin, before treatment with 1ng, 10ng or 100ng/mL TNF (R&D systems, Minneapolis, MN; cat# 410-MT) for 20 mins. Primary CMFs were collected for RNA or protein isolation thereafter.

#### Organoid and Myofibroblasts Co-culture

Briefly, passaged organoids were broken into single cells mechanically by passing through a syringe and needle and were mixed with CMFs at a ratio of 10:1 into Matrigel in a 24-well plate. Co-cultures were grown and maintained in 60% IntestiCult™ (STEMCELL, Seattle, WA; cat# 06005) and 40% CMFs-growth medium at 37°C. The identity and purity of isolated primary CMFs was determined by immunofluorescence for different myofibroblast cell populations (Supplementary Figure 4A).

#### Genotypic Recombination in Co-culture Experiments

Effects of the TNFR1^-/-^ mesenchymal genotype on the epithelium was determined by a series of recombination co-culture experiments in which control or TNFR1^-/-^ organoids were co-cultured with control or TNFR1^-/-^ CMFs for 5 days as described above.

#### Co-culture Experiments with r-RSPO3 and anti-RSPO3 Treatments

Lgr5::EGFP-IRES-CreERT2 organoids co-cultured with TNFR1^-/-^ CMFs were supplemented with increasing doses (5-50 ng/mL) of recombinant-Rspo3 (rRSPO3) (R&D systems, cat# 4120-RS) at passage for 5 days to determine its effect on stem cells, relative to Lgr5::EGFP-IRES-CreERT2 organoids co-cultured with control CMFs. Similarly, Lgr5::EGFP-IRES-CreERT2 organoids co-cultured with control CMFs were treated with an anti-RSPO3 antibody (R&D systems, cat# AF35001) and its IgG isotype control (R&D systems, cat# 5-001-A) in increasing doses of 0.005– 5 µg at passage for 5 days to investigate its inhibition of stem cells, relative to Lgr5::EGFP-IRES-CreERT2 organoids co-cultured TNFR1^-/-^ CMFs. The co-cultures were incubated for 10 min with 2 mg/mL FM4-64X (Thermofisher Scientific, San Diego, CA; cat# F34653) diluted in PBS and were subsequently imaged using a Zeiss LSM 710 Inverted Confocal Microscope (Zeiss, Oberkochen, Germany). Subsequently the co-cultures were processed for RNA isolation for transcript quantification of stem cells.

#### Immunostaining

Immunostaining for paraffin embedded native tissue and organoids as well as whole mounts of co-cultures was carried out as per established protocols. Antibodies used in this study are listed in Supplementary Tables 1 and 2.

#### RNAscope – In-situ hybridization

*In-situ* hybridization (ISH) was performed according to ACD’s manual RNAscope protocol the RNAscope^®^ 2.5 HD Detection Reagents-BROWN (ACD, Newark, CA; cat# 322310). The single-plex RNAscope probes include mouse *Lgr5* (cat# 312171); mouse *Ephb3* (cat# 510251); mouse *Bmp4* (cat# 401301); mouse *Pdgfra* (cat# 480661); mouse *Gli1* (cat# 311001); mouse *Foxl1* (cat# 407401); and mouse *Rspo3* (cat# 402011). The mouse assay positive control included the RNAscope^®^ Control Slide-Mouse 3T3 Cell Pellet (cat# 310023).

#### Immunoblotting

Whole cell lysates from native colonic tissue or primary mesenchymal cells of control or TNFR1^-/-^ animals were prepared as per established methods. Primary and secondary antibodies are listed in Supplementary Table 3. Antibodies were detected using the LI-COR ODYSSEY 9120 infrared imaging system. Densitometric analyses of the band intensities were performed using ImageJ software^34^ (version 1.46; National Institutes of Health, Bethseda, MA). Primary control CMFs were seeded at 1x10^3^ cells/mL in 6-well plates grown to 70% confluency were treated with 10 ng/mL EGF for 5 min and 20 min independently as positive controls for the activation of the PI3k and MAPK pathways. To determine the role for ITGA6 in the regulation of proliferation and/or migration of TNFR1^-/-^ CMFs, a blocking antibody directed against ITGA6 was exposed to TNFR1^-/-^ CMFs grown to 70% confluence (i.e., for 5 d). p27^Kip1^ regulates smooth muscle cells and fibroblasts proliferation and migration synchronously. ^35, 36^ Dose escalation studies of TNFR1^-/-^ CMFs determined that 0.2 µg of anti-ITGA6 restored p27^Kip1^ expression to control levels (Supplementary Figure 8A). In other experiments, control or TNFR1^-/-^ CMFs were seeded at 1x10^3^ cells/mL in 24-well plates and at 70% confluency were treated with 0.2 µg/mL of anti-ITGA6 and IgG isotype control for 5 days for whole cell lysate preparation.

#### Wound Healing Assay

Wounds ∼1 mm in width were applied by scratching confluent monolayers of control or TNFR1^-/-^ primary CMFs with pipette tips were cut obliquely (45° angle) with a sterile razor blade. Cells in suspension were removed by aspiration with PBS. Phase contrast images were taken at 0 h, 2 h, 4 h, 16 h and 24 h, after scratching, using a Leica DM IL LED inverted microscope (Leica, Wetzlar, Germany) equipped with a 5x/0.12 NA objective and a digital camera. Wound closure areas were measured with ImageJ by subtracting the total amount (A) of greyscale pixel counted in the cell-free area remaining after 24 hours from the initial wound area; hence: “Wound closure area” [pixel] =A_initial_ – A_T_ hrs. For antibody blocking experiments, control or TNFR1^-/-^ CMFs were seeded at a density of 1x10^3^ cells/mL in 24-well plates and treated with an anti-ITGA6 antibody and its IgG isotype control at a concentration of 0.2 µg/mL for 0 – 16 h at passage.

#### RNA Isolation, cDNA Synthesis and qRT-PCR

Total RNA from the native tissue or primary CMFs was extracted using the Ambion^®^ PureLink^®^ RNA Isolation Kit (Thermofisher Scientific, cat# 12183025) according to the manufacturer’s protocol. cDNA was synthesized from 50-300 µg of treated RNA for each sample using Verso cDNA Synthesis Kit (Thermofisher Scientific, cat# AB1453B) according to manufacturer’s recommendations using a CFX384 Touch™ Real-Time PCR Detection System (Bio-Rad, Irvine, CA). Quantitative PCR reaction for a subset of primers was performed using the TaqMan Fast Advanced Master Mix (ThermoFisher Scientific, cat# 4444557) on a Bio-Rad iQ5 thermocycler. Primer and probe oligonucleotides were obtained from Integrated DNA Technologies (Newark, NJ). *Lgr5*: (Mm.PT.58.12492947); *Ephb3*: (Mm.PT.58.30165903); *Tnfr2*: (Mm.PT.58.30148877), *Gbp2b*: (Mm.PT.58.13670146), and *Actin*: (Mm.PT.39a.22214843). While other qPCR reactions were carried out in duplicates and contained 5.2 µL SYBR^®^ Premix Ex Taq™ II (Tli RNase H Plus), ROX Plus (Takara, Mountain View, CA; cat# RR82WR), 0.2 µL each of sense and antisense primers (Supplementary Table 4) at 10 µM, 2 µL cDNA or no reaction control with 2.4 µL nuclease free water to give a final volume of 10 µL. Samples were heated to 95⁰C for 3 min followed by 37 cycles of 94⁰C for 15 s, 54⁰C for 20 s, and 72⁰C for 25 s. Primers used in this study are listed in Supplementary Table 4. qRT-PCR data were analyzed by using the comparative ΔΔCt method. ^37^

#### Human Primary Myofibroblast Culture and anti-TNFR1 Experiments

The normal human primary colonic myofibroblast cell line, CCD-18Co (ATCC^®^, Manassas, VA; cat# CRL-1459™) was cultured in Eagle’s Minimum Essential Medium (EMEM) (ATCC^®^, 30-2003™), supplemented with non-essential amino acids, 2 mM L-glutamine, 1 mM sodium pyruvate, and 1500 mg/L sodium bicarbonate and 10% FBS with 1% Penicillin-Streptomycin (10,000 U/mL) and maintained at 37°C in a 5% CO_2_/air mix. The cultures were passaged every 5 days at a ratio of 1:3. To determine the optimal concentration for blocking TNFR1-mediated signaling in human mesenchymal cells, preliminary experiments using human control organoids determined that in response to TNF, epithelial Iĸβα (Supplementary Figures 8B-8D and Supplementary Table 3) is degraded at the highest concentration of 100 ng TNF at 1 h (Supplementary Figures 8B-8D). H-organoids subsequently exposed to increasing doses of H-anti-TNFR1 (Supplementary Table 3) demonstrate that the levels of Iĸβα degradation were restricted in a dose-dependent manner, with little to no effect on the total levels of TNFR1, relative to the IgG isotype control (Supplementary Figures 8B-8D). Therefore, anti-TNFR1 at a concentration of 0.5 µg/mL effectively blocks TNFR1-mediated signaling. Thus, 18Co cells were treated with an anti-TNFR1 antibody and its IgG isotype control at a concentration of 0.5 µg/mL for 5 d at passage. The cells were seeded at a density of 1x10^3^ cells/mL in 24-well plates in parallel for immunostaining and RNA collection.

#### EdU Assay for Proliferation

Whole mounts of organoids or co-cultures were stained with the Click-iT^®^ Plus EdU Alexa Fluor^®^488 Imaging Kit (ThermoFisher Scientific, cat# C10632) as per the manufacturer’s recommendations. For antibody blocking experiments, control or TNFR1^-/-^ CMFs were seeded at a density of 1x10^3^ cells/mL in 24-well plates and treated with an anti-ITGA6 antibody and its IgG isotype control at a concentration of 0.2 µg/mL for 72 h at passage.

### RNA-Seq

Control (TNFR1^+/-^) and TNFR1^-/-^ primary CMFs (passage 6 - 8) were seeded at a density of 1x10^3^ cells/50 µL of Matrigel into 24-well plates and were grown to 70% confluency. At D 5, CMFs were extracted from Matrigel after brief incubation at 4⁰C via centrifugation at 1200rpm for 5 min. Total RNA from primary CMFs was extracted with using the Ambion^®^ PureLink^®^ RNA Isolation Kit (Thermofisher Scientific, cat# 12183025) according to the manufacturer’s protocol. mRNA transcripts from control and TNFR1^-/-^ CMFs were prepared for 2x75 bp sequencing on a HiSeq 4000 (Illumina). Library preparation and sequencing were performed by Children’s Hospital Los Angeles Single Cell, Sequencing, and CyTOF Core Laboratory (Los Angeles, CA). The reads were first mapped to the latest UCSC transcript set using Bowtie2 version 2.1.0 and the gene expression level was estimated using RSEM v1.2.15. TMM method was used for the normalization of the raw count. Differentially expressed genes were identified using the edgeR program. Genes showing altered expression with p < 0.05 and more than 1.5-fold changes were considered differentially expressed. Cluster profiler was used for the GO and pathway enrichment analysis.

### scRNA-Seq

Control (TNFR1^+/-^) and TNFR1^-/-^ primary CMFs were prepared in a similar manner to RNA-Seq. Following centrifugation, cells were reconstituted in 10 mL of HBS/1% BSA solution and were passed through a 40 µm filter to suspend into single cells. To remove dead cells for the final flow, single cell suspensions of control of TNFR1^-/-^ CMFs were stained with the EasySep™ Dead Cell Removal (Annexin V) Kit (STEMCELL, cat# 17899) according to manufacturer’s guidelines. Live cells were quantified and reconstituted in 100-200 µL of HBS/1% BSA solution. Samples with live cell counts of 95% and above only were processed for scRNA-Seq. The transcript abundance measurements were normalized by the per-cell read total, log-transformed, and collapsed to principal components. The top 50 principal components were used for uniform manifold approximation and projection (UMAP). The UMAP was used for clustering. Clustering was performed with the leiden algorithm with a resolution parameter of 1e-5. Markers of each cluster were identified using the Jensen-Shannon specificity score. The significance of each marker was computed via linear regression, which provided an r2 value that could be used to screen the initial gene lists.

For pathway enrichment analysis, the transcript abundances of each cluster were collapsed as a per-cluster, per-gene mean. Genes with an expression level < 0.2 were filtered out. Markers of each cluster were defined as a set of genes whose cluster expression was > 2 times higher than their next highest expression level among the cell clusters. The resulting matrix file was used as input into enrichr. Because clusters 2 and 4 did not exhibit markers of enriched gene expression, these clusters were excluded from enrichr. The search for enriched pathways was run against the KEGG and MSigDB HALLMARK target databases. Pathways with an enrichment adjusted p value (q value) < 0.01 were considered significant and displayed.

