## Supplementary Data for "TNF Receptor 1 regulates colonic mesenchymal cell diversity and the epithelial stem cell niche"

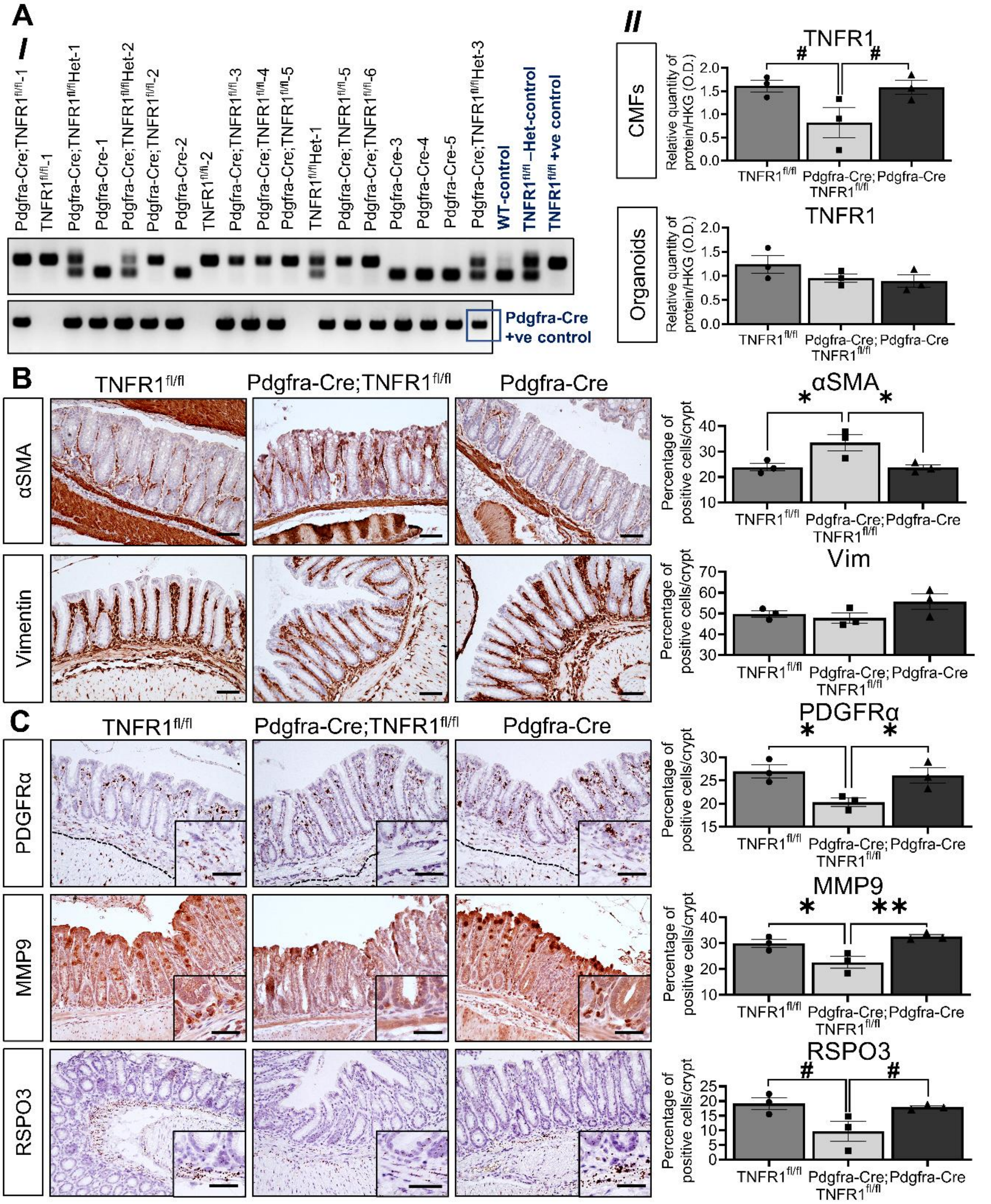


**Supplementary Figure 1 Deletion of TNFR1 in mesenchymal PDGFRα+ cells alters sub-epithelial aSMA+, pericryptal PDGFRα+, Rspo3+ and Mmp9+ mesenchymal cell proportions. Related to Figures 1-3 and 7. (A)** Validation of TNFR1 deletion in the primary colonic mesenchyme of Pdgfra-Cre;TNFR1$^{fl/fl}$ mice via (*I*) genotyping TNFR1$^{fl/fl}$ and Pdgfra-Cre DNA, and (*II*) total TNFR1 protein levels from the primary colonic mesenchymal cells or organoids derived from Pdgfra-Cre;TNFR1$^{fl/fl}$ mice, relative to TNFR1$^{fl/fl}$, and Pdgfra-Cre control animals. Immunolocalization and the percentage of mesenchymal cell markers, **(B)** αSMA and vimentin; **(C)** PDGFRα, RSPO3, and MMP9 in the sub-epithelial and pericryptal niche of distal colon of 6-wk-old Pdgfra-Cre;TNFR1$^{fl/fl}$ mice, relative to TNFR1$^{fl/fl}$ and Pdgfra-Cre controls. The black dotted line indicates the border of the sub-epithelial mesenchymal compartment used for analysis. Subsets represent magnified localization of the markers in the pericryptal niche. Scale bars = 50µm. Error bars are presented as mean ± SEM. Data are analyzed by a one-way Anova, where * $P < 0.05$, and # P = 0.05-0.09. All images are representative of *N* = 3-5 animals.

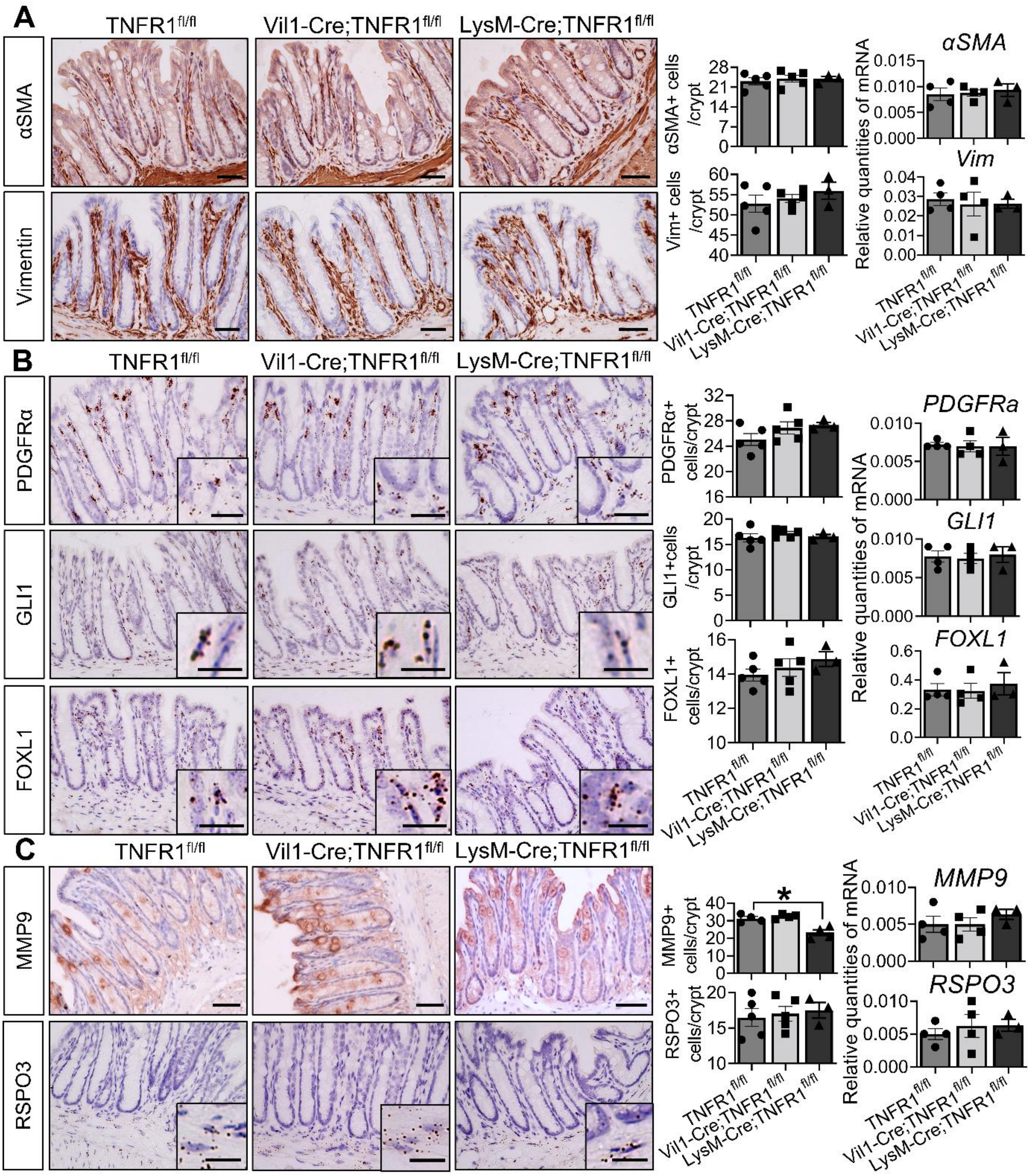


**Supplementary Figure 2 Intestinal epithelial (Vil1-Cre;TNFR1$^{fl/fl}$) and myeloid cell (LysM-Cre;TNFR1$^{fl/fl}$) deletion of TNFR1 does not alter the mRNA expression and numbers of sub-epithelial mesenchymal cells. Related to Figures 1-3.** Immunolocalization, quantification and the transcript levels of mesenchymal cell markers, **(A)** vimentin and αSMA (Scale bars = 50µm); **(B)** PDGFRα (Scale bars = 50µm), GLI1, and FOXL1 (Scale bars = 10µm), **(C)** MMP9 (Scale bars = 50µm), and RSPO3 (Scale bars = 20µm) in the sub-epithelial niche of distal colon of 16w old Vil1-Cre;TNFR1$^{fl/fl}$ and LysM-Cre;TNFR1$^{fl/fl}$ mice, relative to TNFR1$^{fl/fl}$ controls. Subsets represent magnified localization of the markers in the pericryptal niche. Error bars are presented as mean ± SEM. Data are analyzed by a one-way Anova. All images are representative of *N* = 3-5 animals.

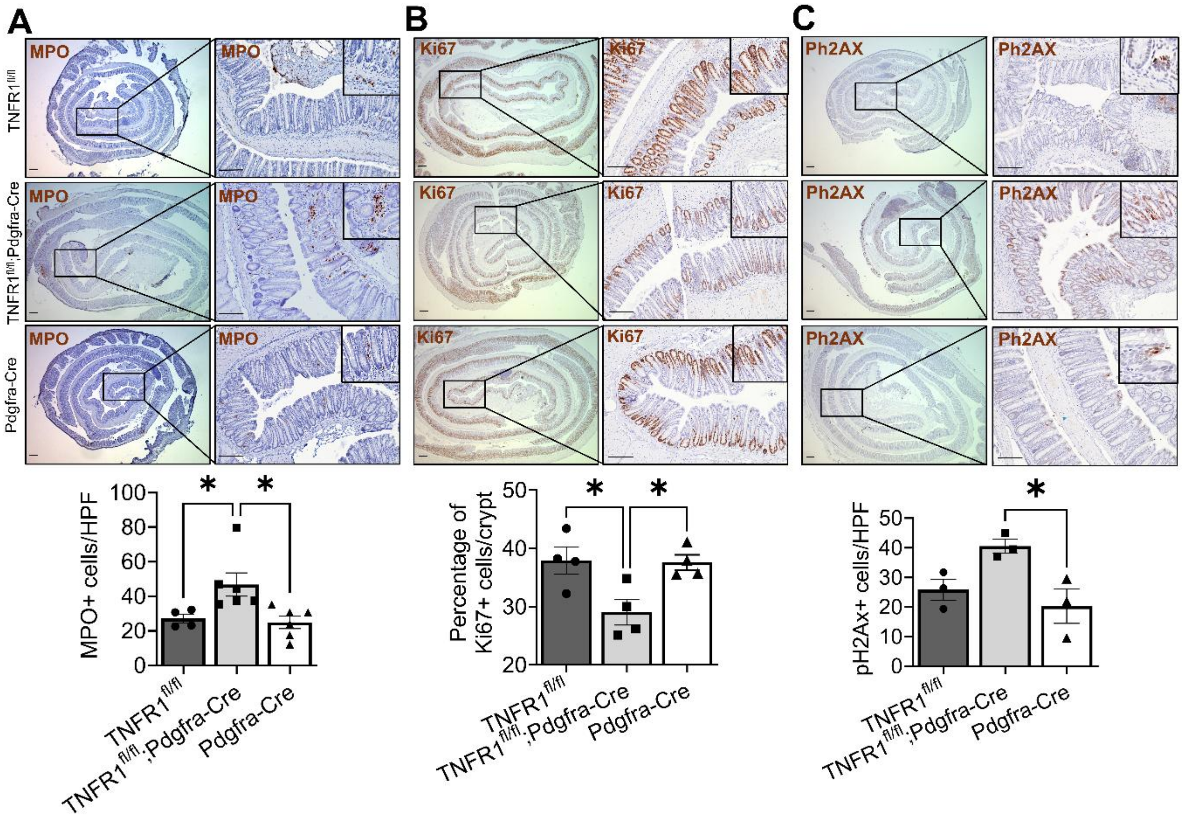


**Supplementary Figure 3 Deletion of TNFR1 in mesenchymal PDGFRα+ cells alters sub-epithelial MPO+, Ki67+, and Ph2AX+ cell proportions. Related to Figures 1-3 and 7.** Immunolocalization and quantification of **(A)** MPO+ cells **(B)** Ki67+ and **(C)** Ph2AX+ cells in the distal colon of Pdgfra-Cre;TNFR1$^{fl/fl}$ mice, relative to TNFR1$^{fl/fl}$, and Pdgfra-Cre control animals (8 wks; *N* = 4-6.) through in situ hybridization. Quantification of MPO+, Ph2AX+ or Ki67+ cells were carried out per High Powered Field (HFP) in crypt epithelial cells and in the sub-epithelial mesenchyme. Scale bars = 200µm. Error bars are presented as mean ± SEM. Data are analyzed by a one-way Anova, where * $P < 0.05$.

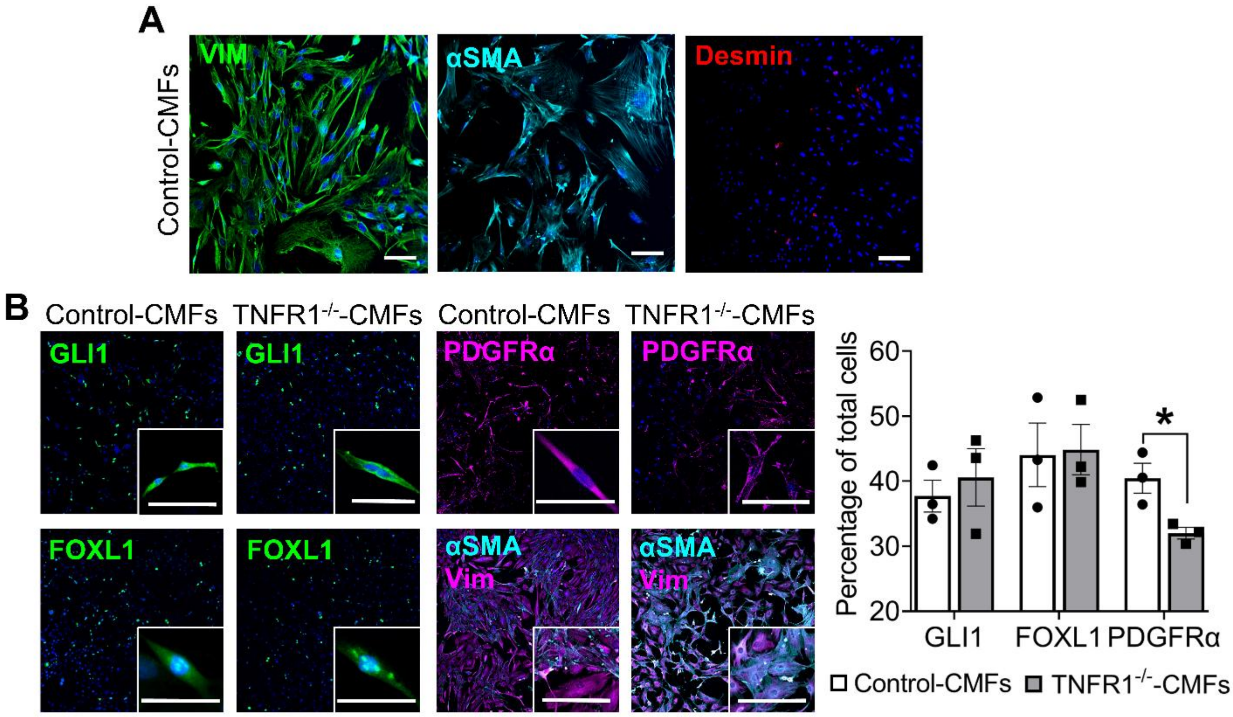


**Supplementary Figure 4 Primary cultures of colonic myofibroblasts express specific markers of the mesenchymal stromal cells. Related to Figures 1-3. (A)** Primary colonic myofibroblasts (CMFs) generated from 16-wk-old control ($TNFR1^{+/-}$) animals, showed a myofibroblast phenotype as demonstrated by positive staining for vimentin and αSMA but negligible staining for desmin by passage 3. **(B)** Immunolocalization and quantification of vimentin, αSMA, FOXL1, PDGFRα, and GLI1 in control ($TNFR1^{+/-}$) or $TNFR1^{-/-}$ CMFs amplified for 5 d in 2D. Scale bars = 25 µm. All images are representative of *N* = 3 animals. Error bars are presented as mean ± SEM. Data are analyzed by an unpaired Student *t* test, where * P < 0.05.

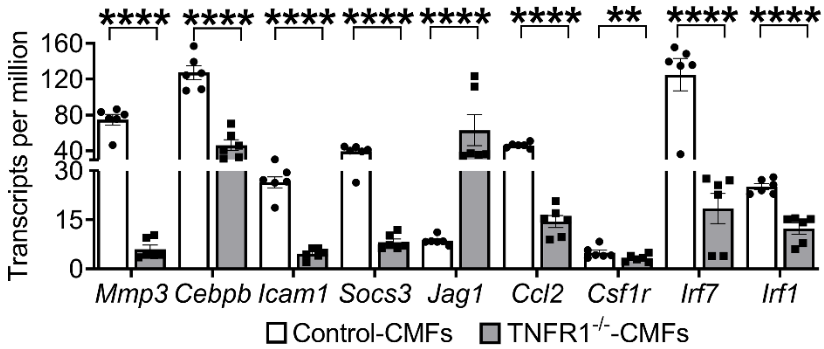


**Supplementary Figure 5 Bulk RNA-Seq profiling of TNFR1-depleted CMFs demonstrates an impairment of TNF pathway genes. Related to Figure 3.** Transcriptome profiling of genes in $TNFR1^{-/-}$ compared to control CMFs identifies alteration in genes in the TNF pathway associated with immunity to microbial infections, mucosal repair, and adaptive immune responses. Values are expressed in transcripts per million (normalized). *N* = 6 animals. Error bars are presented as mean ± SEM. Data are analyzed by an unpaired Student *t* test, where ** P < 0.01, and **** P < 0.0001.

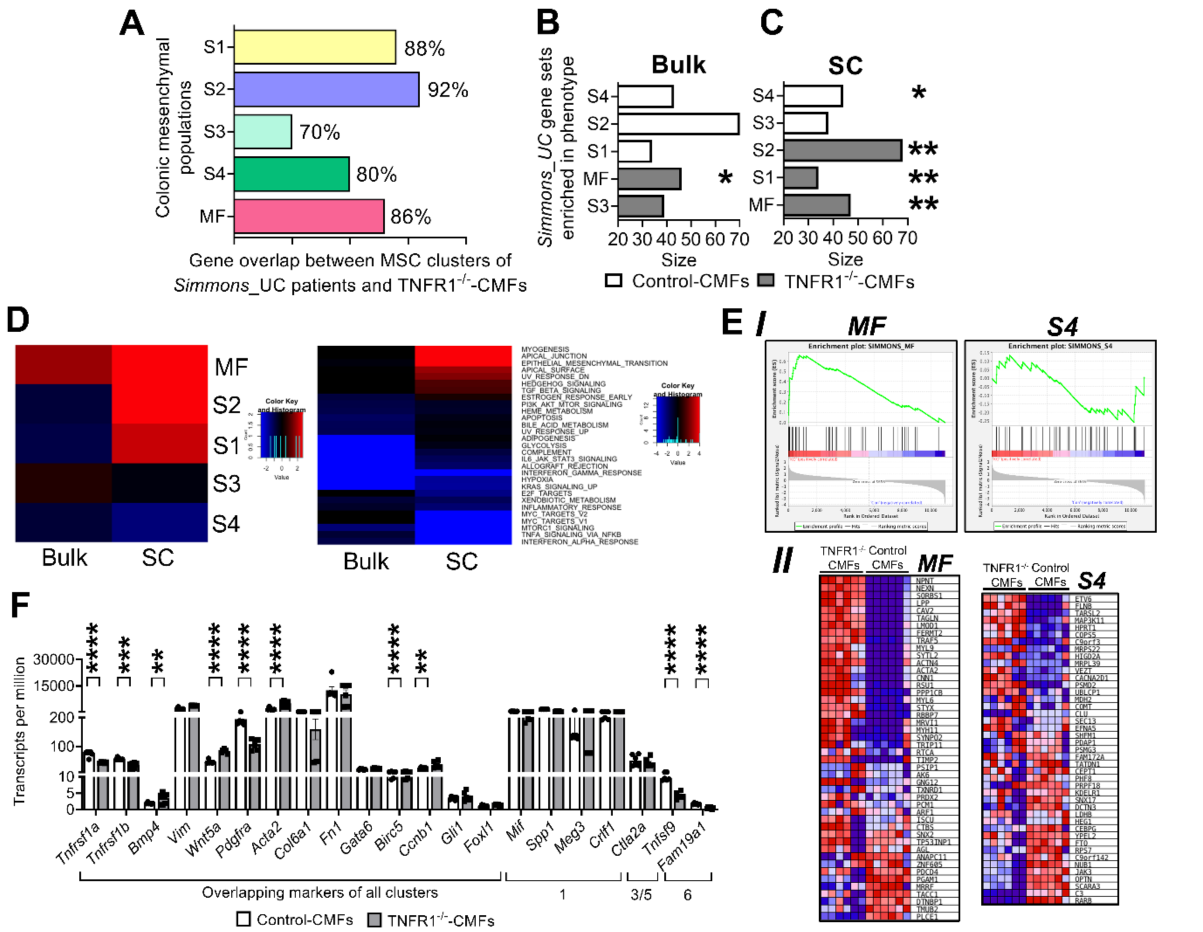


**Supplementary Figure 6 Bulk and single cell transcriptome profiling of primary mouse TNFR1 depleted colonic CMFs show alterations in cytokine signaling pathways of sub-epithelial mesenchymal clusters, similar to the human UC mesenchyme[15]. Related to Figures 2 and 3. (A)** Overlap of genes significantly modified in human UC colonic mesenchymal clusters S1, S2, S3, S4, and MF and in TNFR1$^{-/-}$ CMFs. Mesenchymal gene sets enriched in control and TNFR1$^{-/-}$ CMFs with reference to clusters identified in human UC mesenchyme. Data were analyzed from mouse CMFs **(B)** bulk (*N* = 6) and **(C)** single cell RNA-Seq (*N* = 3) transcriptomes. * P < 0.05; and ** P < 0.01 and human UC mesenchyme. (*Refer to Kinchen and colleagues, Fig. 2, Table S5*[15]) **(D)** Mesenchymal gene clusters and associated pathways of primary control and TNFR1$^{-/-}$ CMFs generated via bulk and single cell RNA-Seq transcriptomes, with reference to the human UC mesenchyme. **(E)** *I* Enrichment plots of mesenchymal clusters, myofibroblasts (MF) and S4 generated by Gene set enrichment analysis (GSEA) showing the 'Hallmark' (Molecular Signatures Database, MSigDB) gene sets enriched from bulk RNA-Seq of mouse colonic control and TNFR1$^{-/-}$ CMFs compared to human UC mesenchyme. The enrichment in the heat maps symbolizing “red” are enriched in TNFR1$^{-/-}$ CMFs and “blue” enriched in control CMFs. The color scale reports the quantitative enrichment score from GSEA. *II* Heatmap of genes encoding the enrichment plot ‘MF’ and ‘S4’ of mouse colonic control and TNFR1$^{-/-}$ CMFs compared to human UC mesenchyme. *N* = 6. **(F)** Bulk RNA-Seq profiling of TNFR1-depleted CMFs demonstrates consistent changes in genes with modified expression patterns identified by single cell RNA-Seq. Error bars are presented as mean ± SEM. *N* = 6. ** P < 0.01; *** P < 0.001; and **** P < 0.0001.

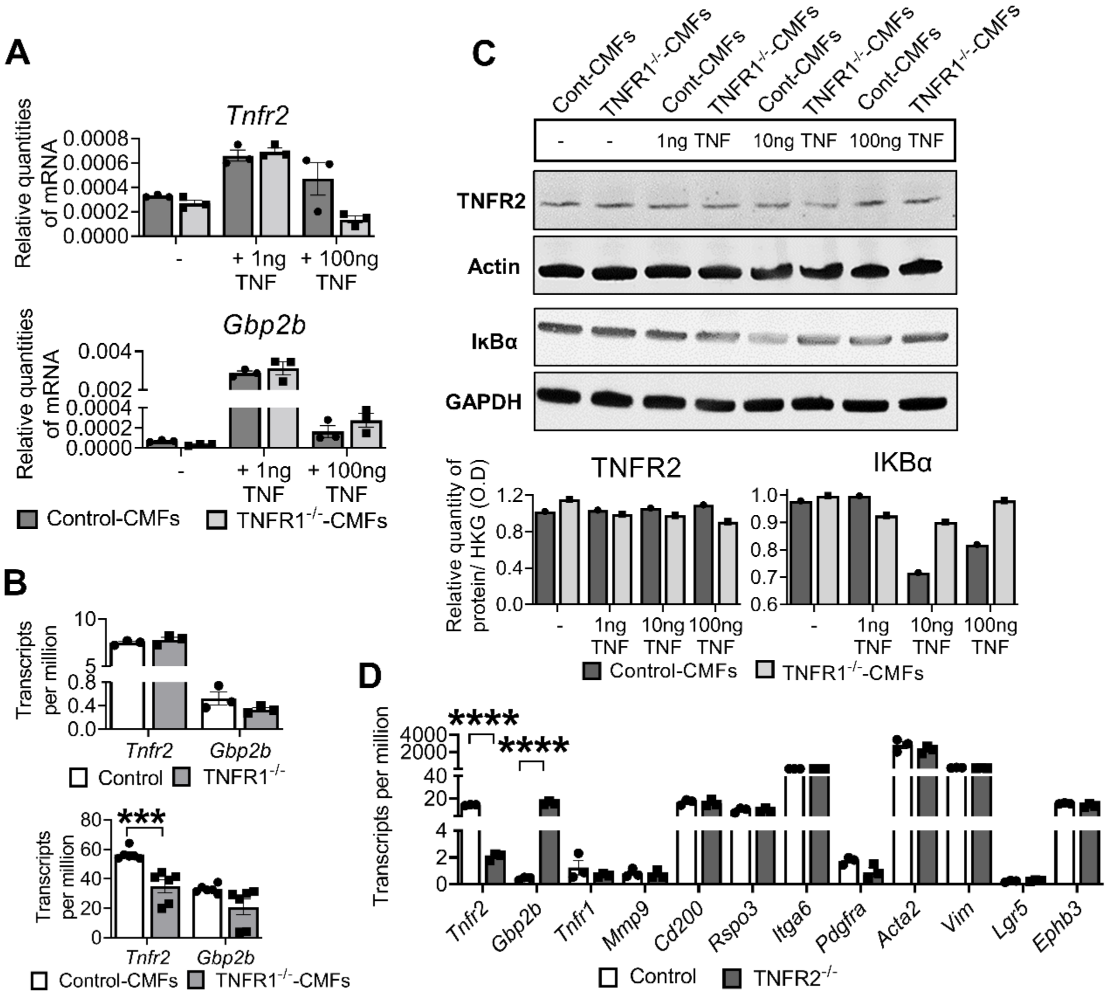


**Supplementary Figure 7 TNFR2-mediated signaling is unaltered in TNFR1 deleted mesenchymal CMFs. Related to Figures 1-3 and 7. (A)** Transcript levels of *TNFR2* and its specific downstream target of a proliferation-associated marker, *Gbp2b*[13] in TNFR1$^{-/-}$-CMFs relative to control-CMFs stimulated with 1ng or 100ng TNF for 20 mins. *N* = 3. **(B)** Transcriptome profiling of genes in the colons of 12-wk old TNFR1$^{-/-}$ mice compared to control (TNFR1$^{+/-}$) mice,[11] or TNFR1$^{-/-}$ and control CMFs, identified no significant changes in the levels of *Tnfr2* and *Gbp2b*. Values are expressed in transcripts per million (normalized). *N* = 3-6 animals. **(C)** Protein levels of TNFR2 and total IκBα in control and TNFR1$^{-/-}$-CMFs stimulated with 1ng, 10ng, or 100ng TNF for 20 mins, quantified relative to the housekeeping genes, actin or GAPDH. *N* = 1. Immunoblots were analyzed via ImageJ. **(D)** Transcriptome profiling of TNFR2 and TNFR1 target genes in in the colons of adult TNFR2$^{-/-}$ mice compared to control (TNFR2$^{+/-}$) mice. *N* = 3 animals. Error bars are presented as mean ± SEM. Data are analyzed by an unpaired Student t test, where *** $P < 0.001$, and **** $P < 0.0001$.

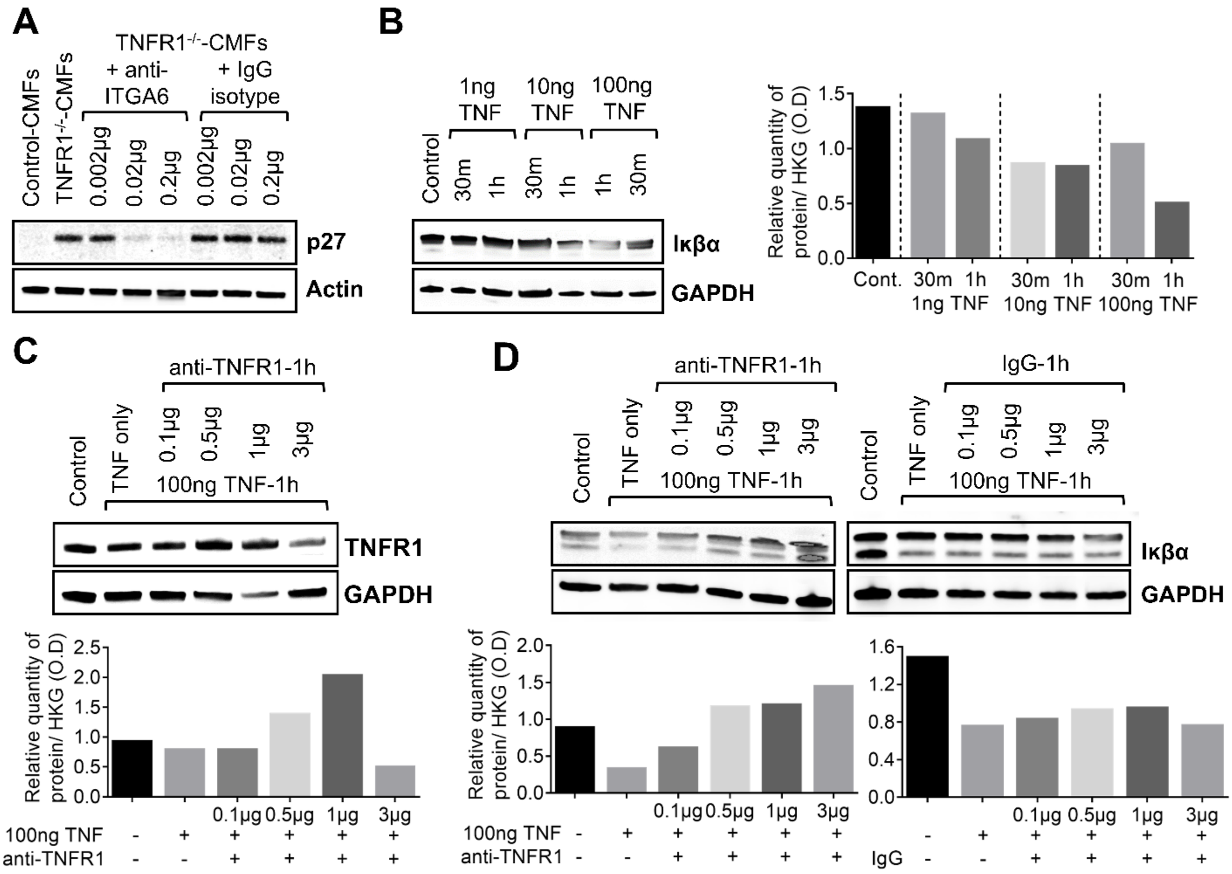


**Supplementary Figure 8 Determination of optimal blocking antibody concentrations for treatments with mouse primary CMFs and human control organoids. Related to Figure 5. (A)** Expression of p27, relative to the housekeeping protein in untreated control and TNFR1$^{-/-}$ CMFs and TNFR1$^{-/-}$ CMFs treated with increasing concentration of anti-ITGA6 antibody and IgG isotype control (0.002 - 2 μg/mL). *N* = 1 sample. Anti-TNFR1 blocks the degradation of Iκβα in a dose-dependent manner. **(B)** Immunoblot of Iκβα levels in H-control organoids treated with increasing concentrations of TNF (1 - 100 ng/mL) for 30m-1h, normalized to GAPDH. Immunoblot of **(C)** TNFR1 **(D)** Iκβα levels in H-control organoids treated with increasing doses of H-anti-TNFR1and its IgG isotype control (0.1 - 3 μg/mL) for an hour preceding treatment with 100ng TNF for 1h. Human colonoids exposed to 0.5 μg of anti-TNFR1 antibody in the presence of 100ng TNF, restores Iκβα levels to control levels. *N* = 1 sample.

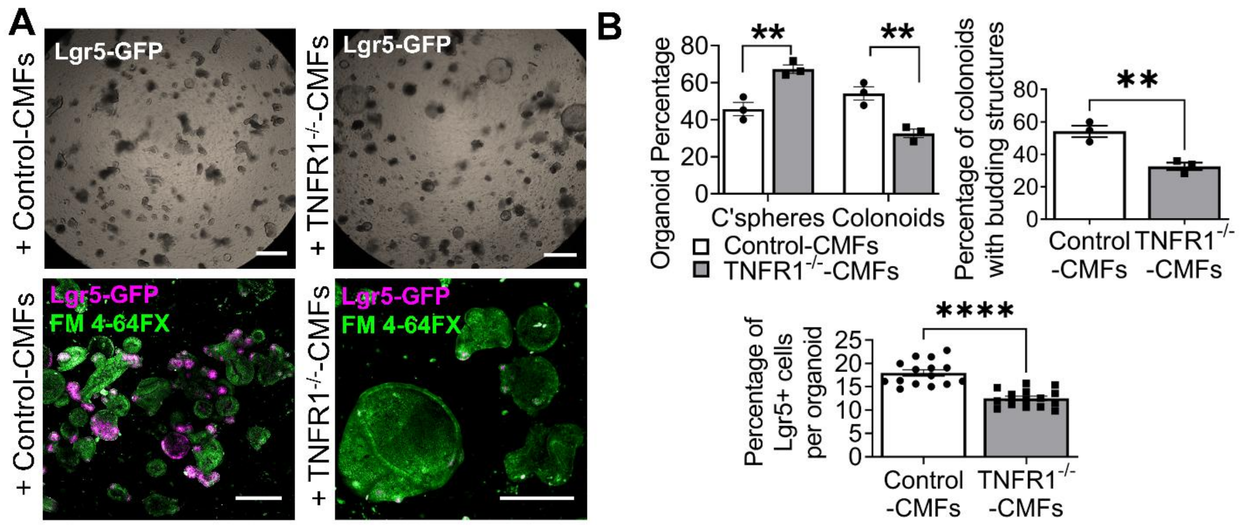


**Supplementary Figure 9 TNFR1 depletion in CMFs downregulates Lgr5+ cells and crypt-like budding structures in Lgr5-GFP colonoids. Related to Figure 7. (A)** Brightfield 2X images (top panel) and 20X (bottom panel) immunofluorescence staining of Lgr5-GFP and FM 4-64X (live-membrane dye) in Lgr5-GFP organoids co-cultured with control or $TNFR1^{-/-}$ CMFs. **(B)** Quantification of the percentage of colonospheres and colonoids, the percentage of colonoids with crypt-like budding structures and quantification of Lgr5+ cells per organoid in co-cultures of Lgr5-GFP organoids and control or $TNFR1^{-/-}$ CMFs. Scale bars = 100 μm. All images are representative of *N* = 3 animals. Error bars are presented as mean ± SEM. Data are analyzed by an unpaired Student *t* test, where ** P < 0.01; and **** P < 0.0001.

**Supplementary Table 1. Primary antibodies for immunofluorescence staining.**

#### Antibodies for immunofluorescence staining

| Antibody (clone) | Affinity | Dilution | Supplier | Catalog |
|---|---|---|---|---|
| Human/Mouse Active Caspase-3 | Rabbit, polyclonal | 1:100 | R&D systems | AF835 |
| E-Cadherin, Clone 34 | Mouse, monoclonal | 1:100 | BD Transduction Laboratories™ | 610404 |
| alpha -Smooth Muscle Actin, Clone 1A4 | Mouse, monoclonal | 1:100 | R&D systems | MAB1420 |
| Anti-Vimentin antibody | Rabbit, polyclonal | 1:100 | Abcam | ab137321 |
| Anti-FOXL1 antibody | Rabbit, polyclonal | 1:50 | Abcam | ab190226 |
| Anti-GLi1 antibody | Rabbit, polyclonal | 1:50 | Rockland Antibodies and Assays | 100-401-223 |
| CD140a (PDGFRA) antibody, Clone APA5 | Mouse, Monoclonal | 1:50 | Thermofisher Scientific | 14-1401-82 |
| Integrin alpha 6 (ITGA6) antibody Clone 6B4 | Mouse, monoclonal | 1:50 | Acris Antibodies | BM6027P |
| Rspo3 antibody | Rabbit, polyclonal | 1:50 | Proteintech | 17193-1-AP |
| Anti-Desmin antibody | Rabbit, polyclonal | 1:100 | Abcam | ab15200 |

**Supplementary Table 2. Primary antibodies for immunohistochemistry staining.**

#### Antibodies for immunohistochemistry staining

| Antibody (clone) | Affinity | Dilution | Supplier | Catalog |
|---|---|---|---|---|
| Anti-MMP9 antibody | Rabbit, polyclonal | 1:100 | Sigma Aldrich | AV33090 |
| CD137 (4-1BB) antibody | Rabbit, polyclonal | 1:100 | Thermofisher Scientific | PA5-116949 |
| CD200 antibody | Rabbit, polyclonal | 1:100 | Bioss Antibodies | BS-6030R |
| TRAF1 antibody | Rabbit, polyclonal | 1:100 | Thermofisher Scientific | BS-1212R |

### Supplementary Table 3. Primary and Secondary antibodies for immunoblotting.

| Antibodies for Immunoblotting and recombinant proteins | | | | |
|---|---|---|---|---|
| **Antibody (clone)** | **Affinity** | **Dilution** | **Supplier** | **Catalog** |
| Anti-Vimentin antibody | Rabbit, polyclonal | 1:1000 | Abcam | ab137321 |
| alpha -Smooth Muscle Actin, Clone 1A4 | Mouse, monoclonal | 1:1000 | R&D systems | MAB1420 |
| Anti-FOXL1 antibody | Rabbit, polyclonal | 1:500 | Abcam | ab190226 |
| Anti-GLi1 antibody | Rabbit, polyclonal | 1:500 | Rockland Antibodies and Assays | 100-401-223 |
| CD140a (PDGFRA) antibody, Clone APA5 | Mouse, Monoclonal | 1:500 | Thermofisher Scientific | 14-1401-82 |
| p27/Kip1 antibody, Clone 225501 | Mouse, Monoclonal | 1:1000 | R&D systems | MAB2256 |
| Phospho-Akt (Ser473) antibody | Polyclonal, Rabbit | 1:1000 | Cell Signaling | 9271 |
| Akt antibody, Clone 40D4 | Mouse, Monoclonal | 1:1000 | Cell Signaling | 2920 |
| Phospho-p44/42 MAPK (Erk1/2) (Thr202/Tyr204), Clone D13.14.4E XP® antibody | Rabbit, Monoclonal | 1:1000 | Cell Signaling | 4370 |
| p44/42 MAPK (Erk1/2) antibody | Rabbit, Polyclonal | 1:1000 | Cell Signaling | 9102 |
| Recombinant Anti-RAP1GAP antibody, Clone Y134 | Rabbit, Monoclonal | 1:1000 | Abcam | ab32373 |
| Anti-RAP1 antibody, Clone 4c8/1 | Mouse, Monoclonal | 1:1000 | Abcam | ab14404 |
| Rspo3 antibody | Rabbit, Polyclonal | 1:500 | Proteintech | 17193-1-AP |
| Anti-β-Actin antibody, Clone AC-15 | Mouse, Monoclonal | 1:5000 | Millipore Sigma | A5441 |
| GAPDH antibody | Rabbit, Polyclonal | 1:5000 | Genetex | GTX100118 |
| TNFRI/TNFRSF1A Antibody | Mouse, Monoclonal | 1:1000 | R&D systems | MAB225 |
| Phospho-IκBα (Ser32) (14D4) Antibody | Rabbit, Monoclonal | 1:1000 | Cell Signaling | 2859 |
| IκBα Antibody | Rabbit, Monoclonal | 1:1000 | Cell Signaling | 9242 |
| IRDye® 800CW IgG Secondary Antibody | Donkey anti-Rabbit | 1:10000 | LICOR | 926-32213 |
| IRDye® 680RD IgG Secondary Antibody | Goat anti-Mouse | 1:10000 | LICOR | 926-68070 |
| Anti-Integrin α6 Antibody, Clone NKI-GoH3 | Rabbit, Monoclonal | 0.2 µg | Millipore Sigma | MAB1378 |
| IgG Control | Rabbit, Polyclonal | 0.2 µg | Millipore Sigma | 12-370 |
| TNFR1 / CD120a / TNFRSF1A Neutralizing Antibody | Rabbit, Monoclonal | 0.5 µg | SinoBiological | 150496-M08H |
| IgG Control | Rabbit, Polyclonal | 0.5 µg | SinoBiological | CR1 |
| R-Spondin 3 Neutralizing Antibody | Sheep, Polyclonal | 0.05 µg | R&D systems | AF35001 |
| IgG Control | Sheep, Polyclonal | 0.05 µg | R&D systems | 5-001-A |
| TNFR2 Recombinant Antibody (ARC0397) | Rabbit, Monoclonal | 1:1000 | Thermofisher Scientific | MA5-35276 |
| Recombinant R-Spondin 3 Protein | CHO-derived mouse protein | 5 ng | R&D systems | 4120-RS |

**Supplementary Table 4. Forward and Reverse primer sequences for SYBR q-PCR reactions.**

| SYBR forward and reverse primers | | |
|---|---|---|
| **Gene** | **Forward** | **Reverse** |
| m-Wnt5A | CAACTGGCAGGACTTTCTCAA | CATCTCCGATGCCGGAACT |
| m-Grem1 | AGACCTGGAGACCCAGAGTA | GTGTATGCGGTGCGATTCAT |
| m-Wnt2b | GGGGATGTTGTCACAGATCA | CTGCTGCTGCTACTCCTGACT |
| m-Bmp4 | TGGGCTGGAATGATTGGATT | CAGTCCCCATGGCAGTAGAAG |
| m-Bmp2 | GGGACCCGCTGTCTTCTAGT | TCAACTCAAATTCGCTGAGGAC |
| m-Rspo3 | TTGACAGTTGCCCAGAAGGG | CTGGCCTCACAGTGTACAATACT |
| m-Fgf-2 | GGCTGCTGGCTTCTAAGTGTG | TTCCGTGACCGGTAAGTATTG |
| m-αSMA | GCATCCACGAAACCACCTA | CACGAGTAACAAATCAAAGC |
| m-Gli1 | AGCCTGAGTCTGTGTATG | CACCACCAGCATGTATTG |
| m-Foxl1 | TGCCGCATTCCACAGCATAGTC | CAAAGTGAGTTCCAGGACAGCCAG |
| m-Vim | CGGCTGCGAGAGAAATTGC | CCACTTTCCGTTCAAGGTCAAG |
| m-Pdgfrα | GCAGTTGCCTTACGACTCCAGA | GGTTTGAGCATCTTCACAGCCAC |
| m-Mmp9 | GCTGACTACGATAAGGACGGCA | TAGTGGTGCAGGCAGAGTAGGA |
| h-β-ACTIN | AGCACGGCATCGTCACCAACT | TGGCTGGGGTGTTGAAGGTCT |
| h-ITGA6 | GAGCTTTTGTGATGGGCGATT | CTCTCCACCAACTTCATAAGGC |
| h-PDGFRα | AACCGTGTATAAGTCAGGGGA | ATTTCTTCCAGCATTGTGAT |
| h-RSPO3 | CAGAATCGCATGACCCAC | AGCAGCCAGAAGCCAGAG |
